# The inhibitory effects of Remodelin on myoblasts differentiation

**DOI:** 10.1101/2024.12.19.629326

**Authors:** Veronica Sian, Andreas Hentschel, Paul E. Görs, Jaakko Sarparanta, Andreas Roos, Per Harald Jonson, Karl W. Smith, Swethaa Natraj Gayathri, Ove Eriksson, Antonello Mai, Dante Rotili, Lucia Altucci, Bjarne Udd, Marco Savarese, Angela Nebbioso

## Abstract

Myoblasts differentiation is a highly regulated and complex process leading to the formation of fused and aligned mature myotubes. Growing interest in the role of epigenetics in muscle differentiation has highlighted epi-modulators as crucial regulators of this process. Our *in vitro* study aimed to explore the potential effects of the inhibition of the acetyltransferase Nat10 on myoblasts differentiation, by using Remodelin, a Nat10 selective inhibitor.

We cultivated and differentiated murine C2C12 myoblasts on ultra-compliant gelatin substrates for up to 16 days and treated them with Remodelin. A combination of morphological analyses, confocal microscopy, transcriptomic profiling (RNA-seq), quantitative proteomics and metabolomics analyses was employed to assess the impact of Nat10 inhibition on myotube formation and maturation. To evaluate the reproducibility of Remodelin effects across myogenic systems and species, L6 rat myoblasts were included as a secondary comparative model.

Remodelin treatment impaired myotube organization, alignment, and structural maturation in both C2C12 and L6 cells compared to untreated controls. In C2C12 cultures, Remodelin also abolished spontaneous myotube contractility. Intersection of transcriptomics and proteomics analyses confirmed that Remodelin effectively slowed myotube formation. Overall, these results indicate that Remodelin broadly affects the regulatory networks involved in skeletal muscle differentiation.

## Introduction

Skeletal muscle is a highly organized and complex tissue, accounting for 45–50% of body mass and performing critical functions such as mobility, metabolism, and thermogenesis (1). The formation, maintenance, and regeneration of skeletal muscle rely on tightly coordinated myogenic programs that integrate extracellular cues with transcriptional and epigenetic regulation (2). During muscle development and regeneration, myogenic precursor cells undergo lineage commitment, cell-cycle exit, and terminal differentiation, processes that are controlled not only by myogenic transcription factors but also by dynamic chromatin remodeling and epigenetic modifications (3–5).

Skeletal muscle exhibits a remarkable ability to adapt and regenerate in response to physiological and pathophysiological stimuli, such as exercise and injury (6). This adaptability depends on the precise regulation of gene expression programs governing myoblast proliferation, differentiation, and myotube maturation (6). Epigenetic mechanisms, including DNA methylation, histone modifications, chromatin accessibility, and non-coding RNAs, play a central role in coordinating these transcriptional transitions by enabling or restricting access or by influencing the stability of myogenic regulators such as MyoD, Myf5, Myogenin, and MRF4 to their target genes (7,8).

Epigenetic regulation, along with genetic predisposition, environmental factors, and lifestyle, plays a pivotal role in age-related diseases and the aging process (9–12). Epigenetic changes can alter the expression of genes essential for maintaining muscle function without changing the gene sequences. Understanding the molecular mechanisms and factors influencing aging and age-related muscle weakness and muscle loss is crucial for developing new pharmaceutical strategies and drug therapies aiming at preserving muscle integrity and performance.

The epidrug Remodelin is a small molecule inhibitor of the N-acetyltransferase 10 (Nat10) activity (13). Remodelin has been extensively studied as a regulator of apoptosis, cell proliferation, invasiveness, and oxidative stress, and it has been shown to sensitize cancer cells to DNA-damaging radiotherapy and chemotherapy (14–17). In lamin A/C-deficient cells, Remodelin has shown promise in correcting defects in the nuclear lamina (18). Additionally, in a mouse model of Hutchinson-Gilford Progeria Syndrome (HGPS), this epi-drug has shown improvements in age-related muscle deterioration and weight loss (18). Recent research has highlighted the upregulation of Nat10 in the skeletal muscles of patients with sepsis, where both *in vitro* and *in vivo* studies have demonstrated that Remodelin counteracts muscle atrophy and weight loss during sepsis by inhibiting the ROS/NLRP3 signalling pathway (19).

Notably, recent studies have identified Nat10 as essential regulator of heart development, heart function, and cardiac remodeling through its RNA-binding and RNA acetylation activities (20,21). Nat10 directly binds and regulates mRNA encoding proteins associated with heart contraction and fatty acid β-oxidation, underscoring its role in coordinating metabolic and structural gene programs in striated muscle tissues (20).

In this study, we used C2C12 murine cells, the most common cellular model to mimic skeletal muscle differentiation *in vitro*, to investigate the effects of Remodelin in muscle differentiation. We differentiated C2C12 myoblasts into myotubes up to 16 days by utilizing a recently described 2D cellular scaffold of gelatin hydrogels (22–24). This innovative system successfully allowed myotube formation as well as contraction and maintained aligned muscle tubes. In parallel, we utilized L6 rat myoblasts as a comparative model to assess the conservation and robustness of Remodelin-mediated epigenetic effects across distinct myogenic cell lines.

## Results

### Remodelin inhibits C2C12 myoblasts viability

We performed MTT assay to evaluate the effects of Remodelin (5, 10, 25 µM) on C2C12 cell viability at different time points (Fig. S1a). The drug showed effects already after 24 h of treatment impairing cell viability by 40 % compared to untreated cells in a non dose-dependent manner. After 48 and 72 h of treatment, 5 µM and 10 µM Remodelin led to 25-30% reduction in cell viability, whereas 25 µM caused a reduction of approximately 35%.

We also investigated the effects of Remodelin on cell cycle progression. Flow cytometry (FACS) analysis showed that, after 8 h of treatment, the epi-drug induced a reduction of G1 phase accompanied by a non-significant increase of S phase (Fig. S1b). After 24 h, Remodelin did not induce any significant modulation of cell cycle.

### Remodelin affects C2C12 myogenesis and myotube twitching

We evaluated Remodelin effects on C2C12 cell differentiation using two concentrations, 5 and 25 μM, with treatments administered every 48 h over a 16-day differentiation period. The concentration of 5 μM did not show any relevant phenotypic effects (Videos S1-S2). In contrast, a Remodelin concentration of 25 μM resulted in impairment of myotube formation and contraction, and, therefore, was used for all the subsequent experiments. Nat10 protein level did not change after Remodelin treatment at day 7 and 16 of differentiation (Fig. S2a).

To investigate the effects of the epi-drug on C2C12 morphology, we captured phase-contrast images at various stages of differentiation (Fig. 1a). Interestingly, while control cells exhibited at day 3 a transition from a round shape to a tubular and elongated shape as a result of cell fusion, Remodelin treatment resulted in a slowdown of this process. At day 7, the untreated cells displayed a highly organized architecture of long and aligned polynuclear tubes, whereas Remodelin altered the fusion and alignment steps inducing an irregular myotube organization. The differentiation index (DI) assessed at day 7 of differentiation showed a 55 % in control cells and 37 % in Remodelin-treated cells (Fig. S2b), indicating that Remodelin markedly impairs myogenic differentiation and myotube organization. The latest stage of differentiation (day 16) showed the same irregular tube shape and alignment, nuclei organization, and impairment of contraction shown already at day 7.

**Figure 1.**
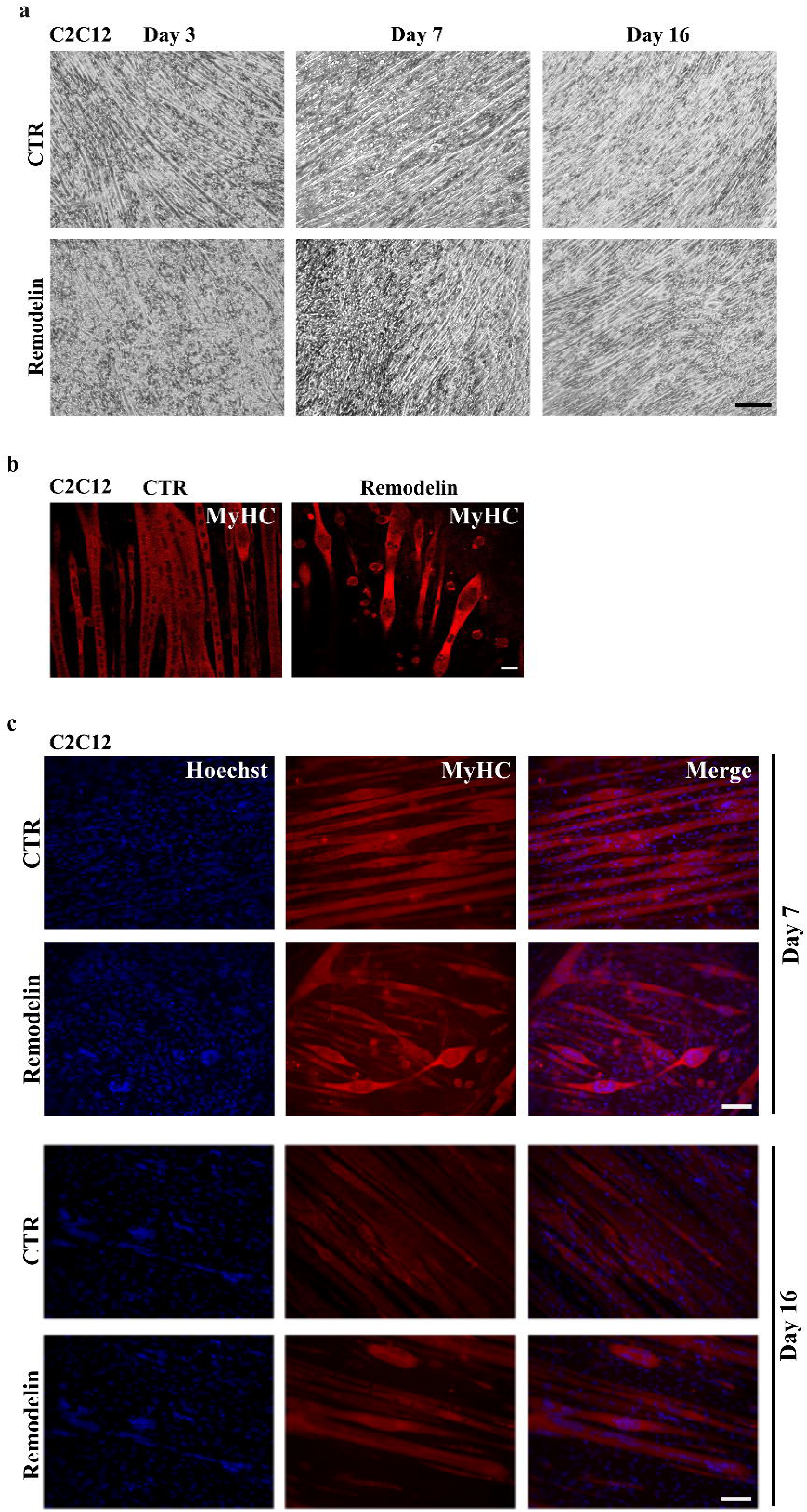
Phenotypic effects of Remodelin on C2C12 differentiation. (a) Representative phase-contrast images of C2C12 cells at three different stages of differentiation (day 3, day 7, and day 16) in control and Remodelin-treated cells. Scale bar: 100 μm. (b) Confocal microscopy images at day 7 of differentiation showing in red Myosin heavy chain II, and in blue nuclei. Scale bar 10 μm. (c) Immunofluorescence analysis of Myosin heavy chain II (red) and nuclei (blue) at day 7 and day 16 of differentiation in control and Remodelin-treated cells. Images were acquired at 20× magnification. Scale bar 100 μm.

Confocal images of myosin heavy chain at day 7 showed that, in the control, nuclei were aligned with each other in long and well-defined myotubes (Fig. 1b). However, the number and the organization of myotubes were impaired in Remodelin-treated cells, and the nuclei did not appear aligned with each other (Fig. 1c).

At day 16 of differentiation, untreated cells showed mature and aligned tubes. In line with previous findings, Remodelin treatment induced the formation of poorly aligned tubes, showing a disorganized nuclei distribution (Fig. 1c).

One notable characteristic of myotube formation is the occurrence of twitching, which is evident already at the intermediate stage of differentiation. At day 7, in the control group, myotubes displayed strong spontaneous twitching confirming myoblasts differentiation into functional myotubes, whereas we did not observe any twitching in Remodelin-treated cells (Videos S3-6). This trend persisted in the late stage of differentiation (day 16), with no twitching detected in Remodelin-treated cells, consistent with our previous observations.

### Omics characterization: transcriptomic and proteomic analyses reveal the downregulation of muscle-related pathways in C2C12

To unveil the effects of Remodelin on C2C12 differentiation, we performed a transcriptomic analysis at days 3, 7, and 16 with and without Remodelin treatment. Two-dimensions Principal Component Analysis (PCA) plot revealed a noticeable difference between control and treatment at day 16. Control cells segregated into three distinct clusters corresponding to day 3, 7, and 16, reflecting the progressive transition from myoblasts to myotubes, while Remodelin-treated cells clustered together at day 7 and 16, indicating a clear defect in maturation (Fig. 2a).

**Figure 2.**
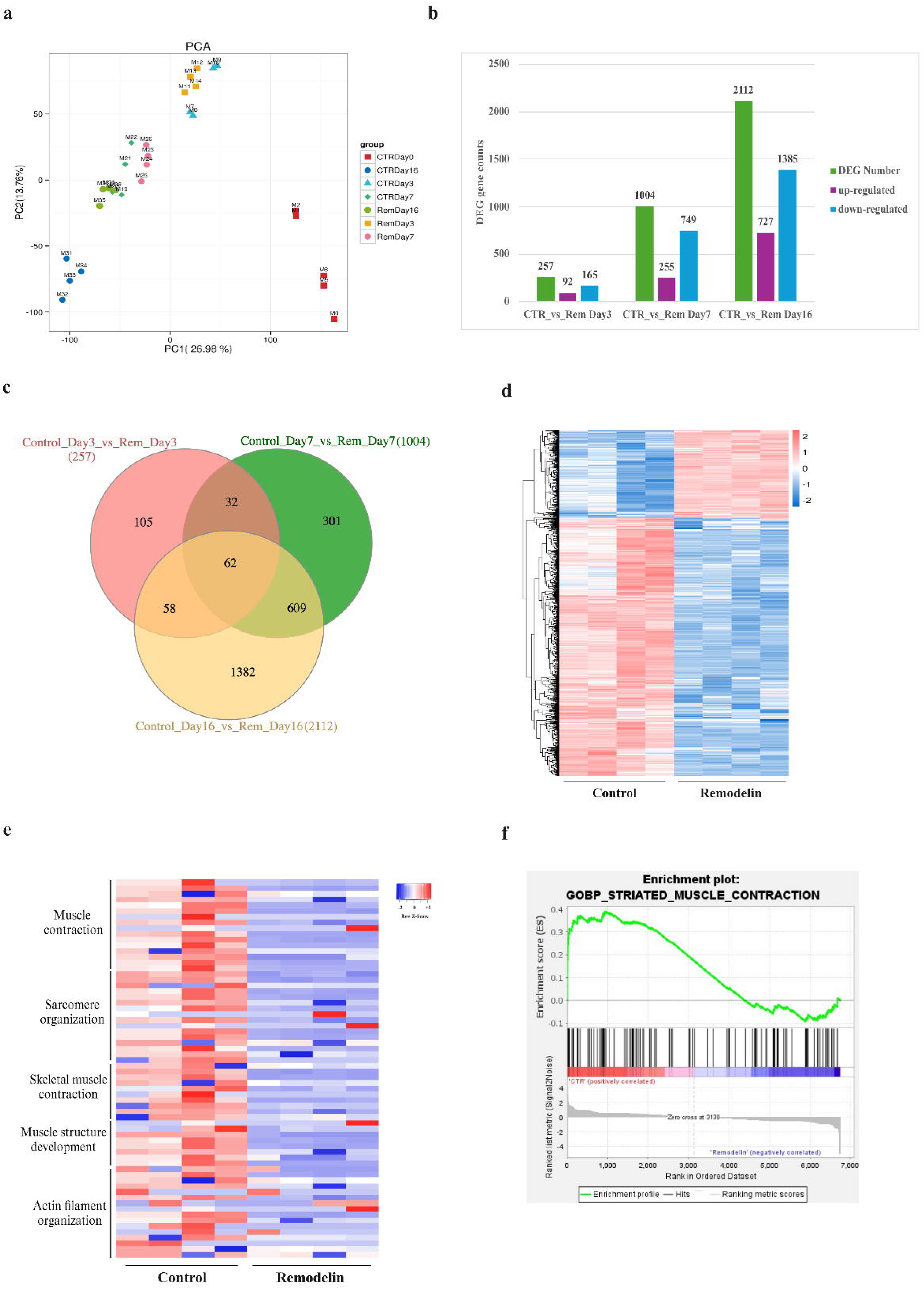
RNAseq analysis of Remodelin-treated C2C12 cells during differentiation. (a) Two-dimensional principal component analysis (PCA) of control and Remodelin-treated C2C12 samples projected along PC1 (26.98%) and PC2 (13.76%), illustrating the progressive separation of samples according to treatment and differentiation stage. (b) Bar plot summarizing the number of differentially expressed genes (DEGs) at day 3, day 7, and day 16 of differentiation. Green bars indicate the total number of DEGs, while purple and blue bars represent up-regulated and down-regulated genes, respectively, following Remodelin treatment. The y-axis denotes DEG counts. (c) Venn diagram representing the shared up-regulated and down-regulated genes among different days of differentiation (day 3, day 7, and day 16). A total of 257, 1004, and 2112 genes were differentially expressed at day 3, 7, and 16 respectively. (d) Heatmap representation of gene expression differences between control and Remodelin-treated cells at day 7 of differentiation. (e) Heatmap of the selected DEGs associated with key myogenic processes, including muscle contraction, sarcomere organization, skeletal muscle contraction, muscle structure development, and actin filament organization. Red and blue color gradients indicate increased and decreased gene expression, respectively. The x-axis represents the different samples of the two compared groups, the y-axis shows individual samples from each experimental group, and the y-axis lists the differentially expressed genes included in the analysis. (f) Gene set enrichment analysis (GSEA) comparing control and Remodelin-treated cells at day 7 of differentiation. The enrichment score (ES) is shown on the y-axis, while the x-axis represents the ranked gene list; vertical black lines indicate the positions of genes belonging to the enriched gene set. Statistical significance was determined using a false discovery rate (FDR) threshold of < 0.05.

Pairwise comparison between controls and the corresponding Remodelin treatment at day 3, 7 and 16 showed a total of 257 (92 up- and 165 down-regulated), 1004 (255 up- and 749 down-regulated), and 2112 (727 up- and 1385 down-regulated) differentially expressed genes, respectively (Fig. 2b). Venn diagram showed the shared dysregulated genes among the different time points. A total of 62 genes were shared among the 3 time points, while 609 genes were shared among day 7 and day 16 of differentiation (Figure 2c). The relatively high number of Differentially Expressed Genes (DEGs) observed on day 16 arises from comparing untreated cells at their mature stage of differentiation with Remodelin-treated cells that have not reached the same level of differentiation.

According to the phenotypic data of myotube formation and contraction, day 7 of differentiation emerged as a crucial point to better dissect the effects of Remodelin in myotube formation. The cluster heatmap visually illustrates the distinction in gene expression levels of 1004 genes between the two groups (Fig. 2d).

We performed Gene Ontology (GO) analysis to determine the potential role of dysregulated genes (Table 1). Enrichment analysis assigned genes to three functional categories: (i) biological process (BP), (ii) cellular component (CC), and (iii) molecular function (MF), and ranked them based on false discovery rate (FDR). The top 10 downregulated of the three categories were associated to skeletal muscle development and maturation, myotube structure and the components of myofibers, and functions of mature fibers. This data corroborated the previously described phenotypic findings, supporting the idea that Remodelin causes a slowdown in myotube maturation and impairs contraction.

**Table 1.**
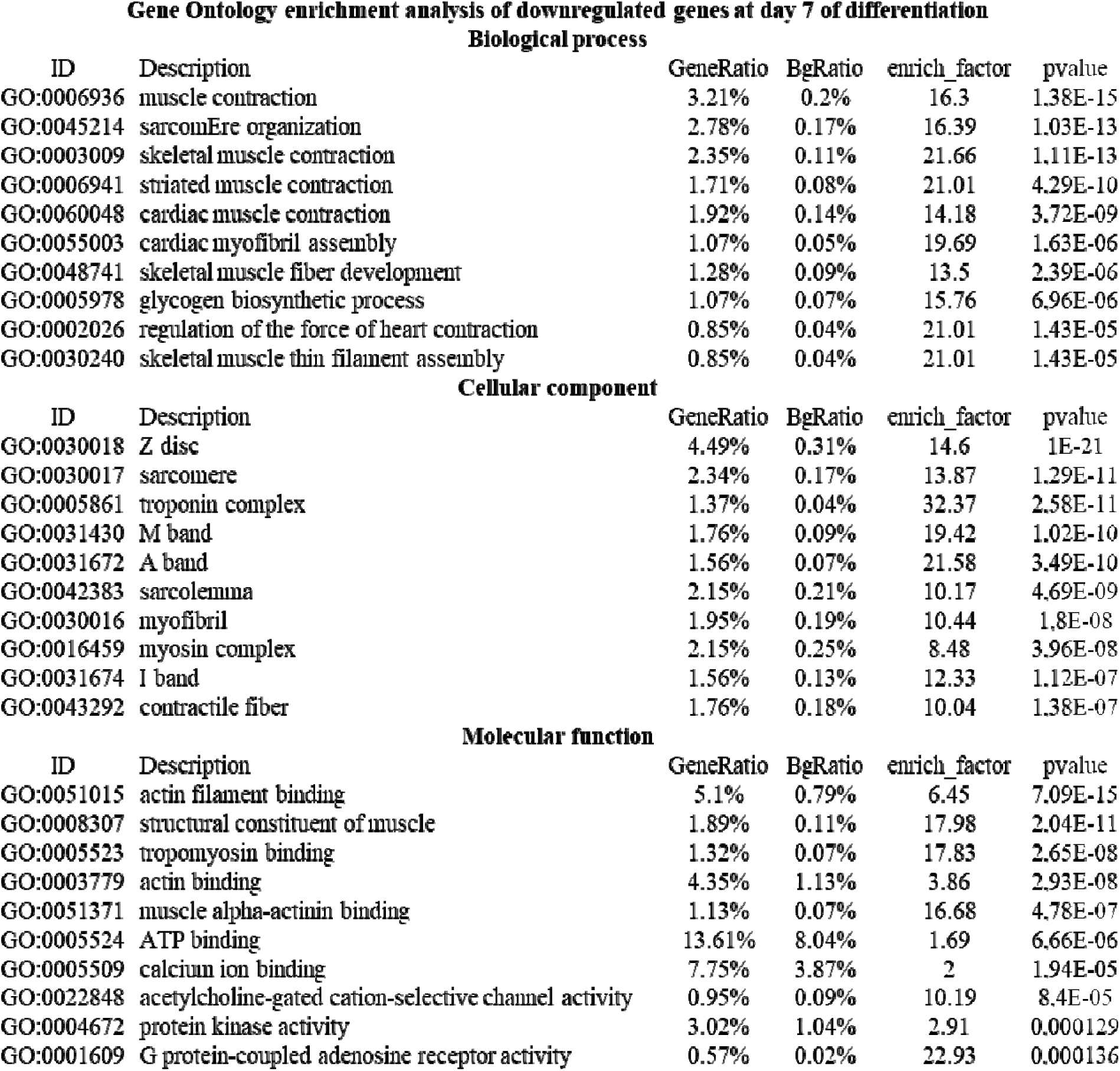
Gene Ontology (GO) enrichment analysis of down-regulated genes following Remodelin treatment in C2C12 cells after 7 days of differentiation. The top 10 enriched terms for biological processes, cellular components, and molecular functions are shown. Terms within each category are ranked according to adjusted p-values.

GO analysis at day 3 (Table S2) and at day 16 (Table S3) of differentiation confirmed the effects of Remodelin also at the early and late stage of myotube formation, corroborating the previous phenotypic findings.

We performed hierarchical cluster analysis on the DEGs involved in significant selected biological processes at day 7 of differentiation (Table S4). Notably, DEGs of muscle contraction, sarcomere organization, skeletal muscle contraction, muscle structure development, and actin filament organization were deregulated. Heatmap analysis revealed a clear grouping according to controls and treatments. All the DEGs involved in the most inhibited pathways strongly decreased upon Remodelin treatment, confirming the effects of the drug in the modulation of muscle differentiation (Fig. 2e). We performed gene set enrichment analysis (GSEA) on the differentially expressed genes among the two groups at day 7 to identify dysregulated signaling pathways. As expected, we found the GOBP_STRIATED_MUSCLE_CONTRACTION gene set correlated with Remodelin treatment (p-value < 0.02) (Fig. 2f).

Mass spectrometry-based proteomic data on untreated and Remodelin-treated cells at day 7 of differentiation unveiled that, out of 3074 quantified proteins, 26 proteins were statistically significantly up-regulated, and 37 proteins were statistically significantly down-regulated upon Remodelin treatment compared to the control group, as shown in the volcano plot (Fig. 3a). In addition, we identified 17 proteins only in the control group and not in the Remodelin group. A heatmap of the proteomic dataset at day 7 of differentiation illustrates the global protein expression patterns in control and Remodelin-treated cells. Clustering analysis revealed a clear separation between the two conditions (Fig. 3b).

**Figure 3.**
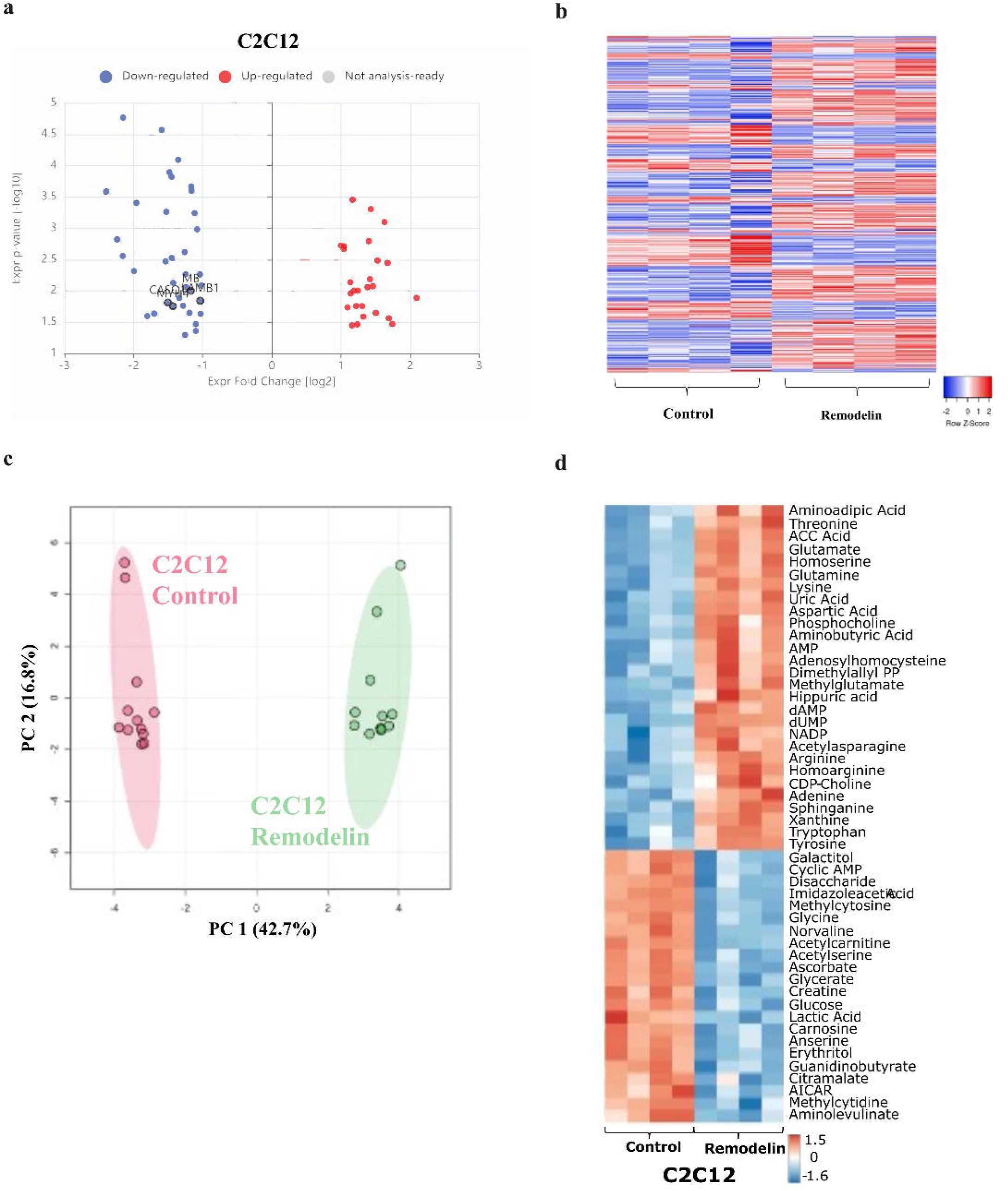
Proteomic and metabolomic profiling of C2C12 cells at day 7 of differentiation after Remodelin treatment. (a) Volcano plot representing the differential protein expression (DEPs) analysis at day 7 in control and Remodelin groups. The number of up- and down-regulated proteins were distinctly identified based on the log-fold change criteria between the two groups, and the values were set between 0.5 for the down-regulated proteins and 2 for the up-regulated. In blue the down-regulated and in red the up-regulated proteins. (b) Heatmap representing relative protein abundance in control and Remodelin-treated cells at day 7 of differentiation. Data are hierarchically clustered based on similarity across samples. Color in each tile represents the scaled abundance value. (c) Principal component analysis (PCA) of metabolomic data showing distinct clustering of control (CTR) and Remodelin-treated samples along PC1 (42.7%) and PC2 (16.8%), indicating treatment-associated metabolic changes. (d) Metabolite profiling is exhibited in a heatmap of 50 most significant annotated metabolites in control and Remodelin-treated samples.

The gProfiler tool revealed the enrichment of biological process in both downregulated (Fig. S3a) and up-regulated (Fig. S3b) proteins. The analysis of the down-regulated terms revealed that Remodelin impaired the calcium ion binding, extracellular matrix structural constituent, muscle cell differentiation, muscle structure development, laminin-1 complex (Fig. S3a). The protein-protein interaction (PPI) network was constructed using STRING database for the dysregulated proteins (Fig. S4a-b). STRING protein-protein interactions identified 36 nodes and 70 edges, with a PPI enrichment p-value < 1.0×10^−16^, and the analysis revealed six different interactions (Fig S4a) for the down-regulated proteins. We performed the GO analysis using the GOrilla analysis tool for the down-regulated proteins (Fig. S4c). Once more, the extracellular matrix constituent and the calcium ion binding were enriched in the Remodelin group compared to the control.

The gProfiler analysis of the up-regulated proteins revealed the alteration of pathways related to metabolic processes and mitochondria (Fig. S3b). STRING analysis revealed 26 nodes and 13 edges, with a PPI enrichment p-value 5.6×10^−05^ (Fig. S4b).

Combined analysis of the transcriptomics and proteomics results identified 2 up-regulated terms in common and 7 down-regulated terms (Fig. S3c). The two common up-regulated genes are the Aldehyde Dehydrogenase 3 Family Member A1 (*Aldh3a1*) and NAD(P)H quinone oxidoreductase 1 (*Nqo1*). The down-regulated genes are the Heat shock protein beta 7 (*Hspb7*), Myoglobin (*Mb*), Myosin heavy chain 4 (*Myh4*), Insulin-like growth factor-binding protein 5 (*Igfbp5*), Calsequestrin 1 (*Casq1*), and N-myc downstream-regulated gene 2 (*Ndrg2*).

### Metabolomic profile of Remodelin effects on C2C12 myotube maturation

To further analyze the effects of Remodelin on C2C12 differentiation, we performed metabolomic profiling at day 7. PCA revealed distinct clustering of control and Remodelin-treated samples, separated along PC1 (42.7%) and PC2 (16.8%) (Fig. 3c). This clusterization is consistent with the previous omics analyses and the previous phenotypic results. Heatmap representation of the 50 most significant metabolites highlighted specific changes in key metabolic pathways (Fig 3d). Selected metabolites demonstrated that Remodelin treatment was associated with altered levels of glucose, uric acid, NADPH, UDP-glucuronic acid, acetylcarnitine, and lactate, as illustrated by boxplots (Fig. S5). These results suggest that Remodelin impacts both energy metabolism and redox balance, potentially contributing to its modulatory effects at the cellular level.

### Remodelin affects histone acetylation during differentiation in C2C12

We characterized the epigenetic landscape of C2C12 cells during differentiation through Western blot analyses at three different stages, day 3, day 7, and day 16 (Fig. 4a-b). Our findings indicated that histone acetylation level increased at day 7 and day 16 during myotube maturation. Specifically, histone H3K18ac exhibited an upregulation at day 7, while histone H4K8ac and histone H3K19/14ac at both day 7 and day 16. However, the differentiation did not influence histone methylation levels (Fig. 4b).

**Figure 4.**
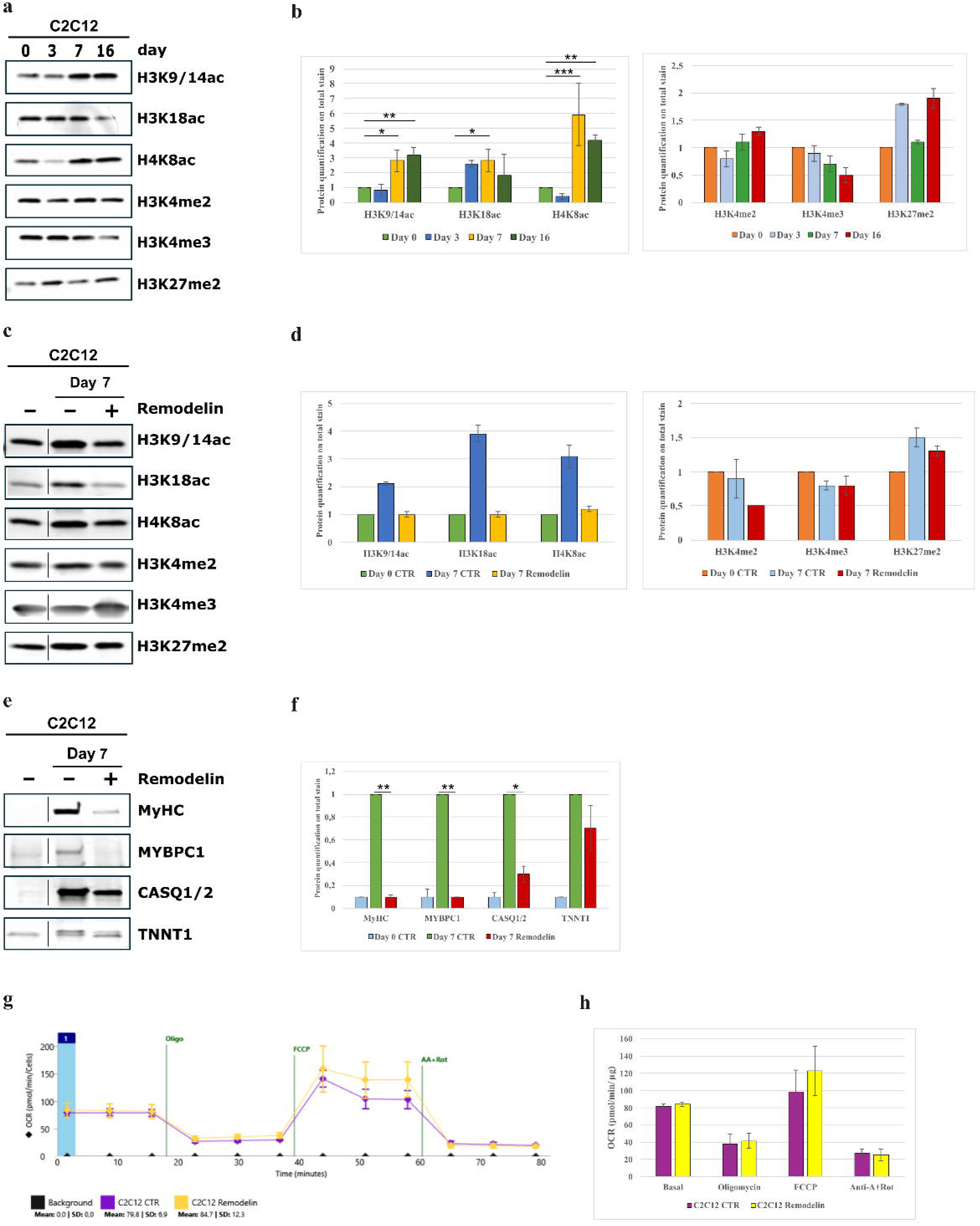
Western blot analysis of epigenetic modulation and myogenic differentiation markers in C2C12. (a) Representative Western blots showing histone acetylation marks (H3K9/14ac, H3K18ac, H4K8ac), and histone methylation marks (H3K4me2, H3K4me3, H3K27me2) in control C2C12 cells at different stages of differentiation. (b) Histograms representing histone quantification normalized on total protein stain. Protein levels are expressed relative to control cells at day 0. Data are presented as mean ± SEM (n = 3). Statistical significance is indicated as follows: * p ≤ 0.05, ** p ≤ 0.01, *** p ≤ 0.001. (c) Representative Western blots illustrating changes in histone acetylation and methylation in C2C12 cells during differentiation following Remodelin treatment. (d) Quantification of epigenetic markers normalized to total protein staining and expressed relative to control cells at day 0. (e) Representative western blots of differentiation markers at day 7 of differentiation. Myosin heavy chain II (MyHC), Mybpc1, Calsequestrin 1/2 (CASQ1/2), and Troponin T (TNNT1) in control and Remodelin-treated cells. (f) Bar chart representing the quantification of proteins, normalized to the total protein stain. Control day 7 was considered as reference control. Data are representative of three independent experiments. Statistical significance is indicated as follows: * p ≤ 0.05, ** p ≤ 0.01. (g) Measurements of oxygen consumption rate (OCR) before (basal) and after addition of oligomycin, FCCP, and antimycin A in combination with rotenone, in C2C12 myoblasts after 24 h of Remodelin 25 μM treatment. (h) Averaged OCR values of two biological experiments. Data are presented as mean±SD (n=20 for control and n=30 for Remodelin samples).

We characterized the effects of Remodelin on epi-modulation after 7 days of differentiation. Western blot analyses revealed that, as expected, Remodelin induced downregulation of histone acetylation compared to the respective controls, and it had no effects on histone methylation status (Fig. 4c-d).

### Remodelin modulates markers of differentiation in C2C12

To confirm the omics findings on the impact of Remodelin in C2C12 differentiation and myotube formation, we performed Western blot analyses of specific protein markers of differentiation at day 7 (Fig. 4e-f). Remodelin-treated cells exhibited a significant reduction in the protein levels of MyHC, Mybpc1, CASQ1/2, and Troponin T compared with control cells (Fig. 4e–f). Conversely, robust upregulation of these markers was observed in controls at day 7 compared to the control at day 0, consistent with efficient myotube formation and maturation.

### Mitostress test

To assess the possible effects of Remodelin on mitochondrial function, we performed a mitostress test using the Seahorse XF Pro Analyzer. C2C12 myoblasts were treated with Remodelin (25 µM) for 24 h prior the assay. After sequential addition of oligomycin, FCCP, and antimycin A in combination with rotenone, oxygen consumption rate (OCR) parameters were assessed. Remodelin treatment did not significantly alter basal respiration, proton leak, or non-mitochondrial respiration in C2C12 cells after 24 h of treatment (Fig. 4g-h).

### Nat10 expression in C2C12 and L6

We also examined the effects of Remodelin in a second myogenic model, the L6 rat myoblasts. These cells displayed higher Nat10 expression at the myoblast stage compared with C2C12 cells (Fig. 5a). Immunofluorescence confirmed Nat10 expression and nuclear localization in both C2C12 and L6 myoblasts (Fig. 5b). During differentiation, Nat10 expression followed a peculiar pattern in C2C12 cells: its levels decreased at day 3 and 7 of differentiation followed by an increase at day 16, corresponding to mature myotubes (Fig. 5c). A similar trend was observed in L6 cells, where Nat10 protein level showed a significant reduction already at day 3 (Fig. 5d).

**Figure 5.**
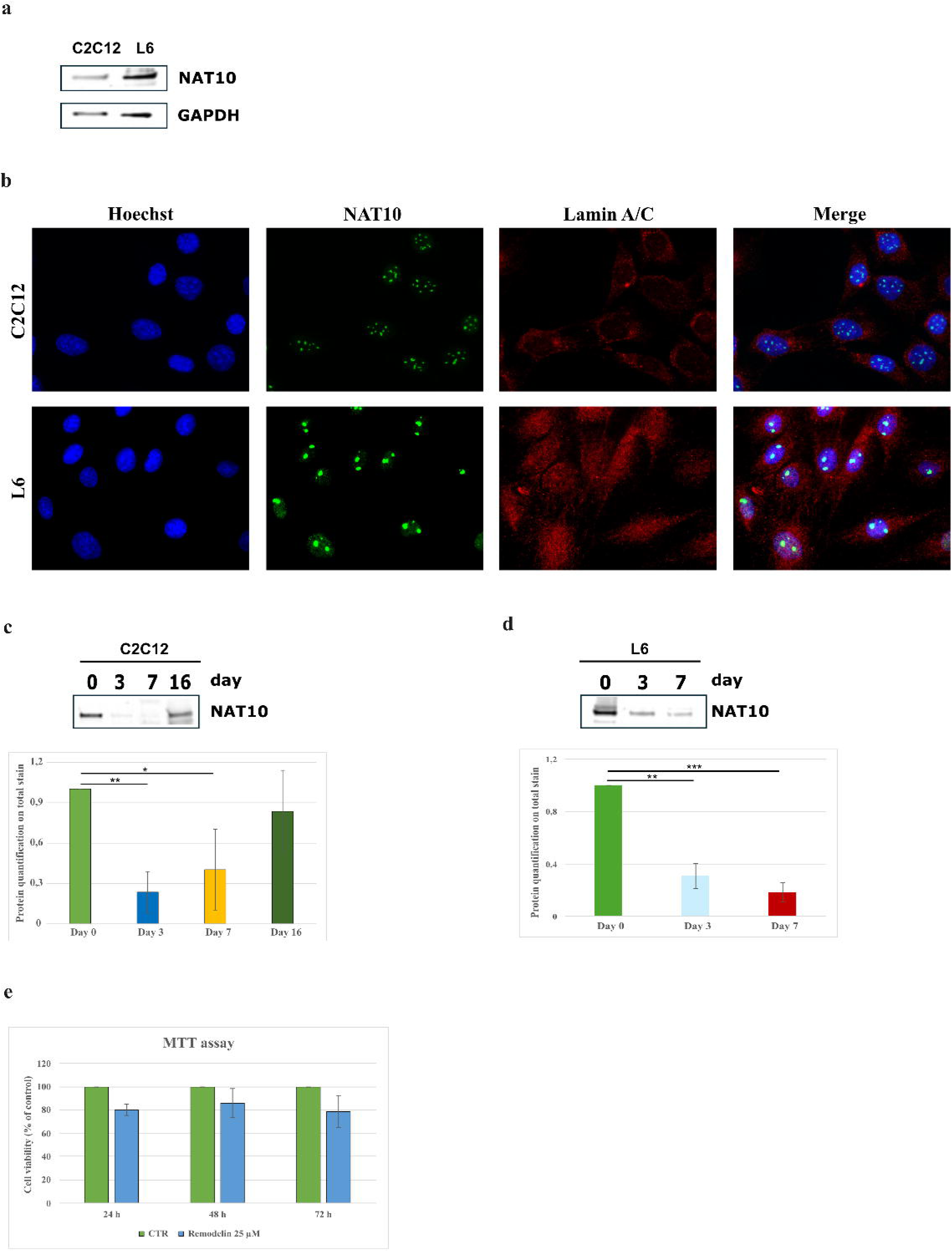
NAT10 expression in C2C12 and L6. (a) Representative Western blots of NAT10 protein level in C2C12 and L6 myoblasts. GAPDH was used as loading control. (b) Immunofluorescence analysis of NAT10 expression and localization in C2C12 and L6 myoblasts. Nuclei are shown in blue, NAT10 in green, and Lamin A/C in red. (c) Representative Western blots and quantification of NAT10 expression in C2C12 cells at day 0, day 3, day 7, and day 16 of differentiation. (d) NAT10 expression and its quantification in L6 differentiation at day 0, 3, and 7. (e) MTT assay showing the percentage of L6 cell viability after 24, 48, and 72 h of treatment with Remodelin (25 µM).

MTT assay showed that Remodelin 25 µM led to 20 % reduction of L6 cell viability after 24 h of treatment, 15% after 48 h, and 20 % after 72 h (Fig. 5e).

### Effects of Remodelin on L6 differentiation

L6 myoblasts were initially seeded onto the gelatin hydrogel scaffold used for C2C12 differentiation; however, unlike C2C12 cells, L6 cells failed to adhere to the hydrogel, preventing their use in this system. Thus, L6 cells were differentiated on collagen-coated dishes. Although L6 cells initially differentiate efficiently, a decline in myogenic marker expression has been reported after approximately 6 days in culture (25), restricting the use of the model to early stages of myogenesis.

L6 cells were therefore differentiated for up to 7 days in DMO medium with Remodelin treatments (25 µM) administered every 48 h. Phase-contrast microscopy revealed myotube formation in both control and Remodelin-treated cultures; however, L6 myotubes appeared larger and less aligned compared with C2C12 myotubes as early as day 3 of differentiation. Notably, Remodelin treatment further delayed myogenic progression, resulting in fewer and less organized myotubes (Fig. S6a). Immunofluorescence analyses corroborated these observations, showing pronounced alterations in myotube morphology and disrupted nuclear organization in Remodelin-treated L6 cells relative to controls at day 7 of differentiation (Fig. S6b).

### Omics insights into L6 myotube formation

RNAseq analysis was performed on L6 cells at day 3 and day 7 of differentiation. PCA analysis showed that the control and Remodelin samples at day 3 and day 7 at both time points clustered by condition rather than by differentiation stage, supporting the decision to perform differential expression analysis on the combined datasets (Fig 6a). Remodelin treatment resulted in 946 up-regulated and 1,771 down-regulated genes. The corresponding heatmap of differentially expressed genes illustrates distinct expression patterns between the two groups (Fig 6b). gProfiler analysis revealed that the GO enriched biological process (BP) terms were linked to cytoskeleton organization and myotube differentiation, while the cellular component (CC) terms included cytoskeleton, myofibril, and sarcomere. The most significantly down-regulated molecular function (MF) terms were associated with tubulin, actin and laminin binding, and extracellular matrix binding (Table S5). The GO BP terms for the up-regulated genes were predominantly associated with ribosomal biogenesis, rRNA metabolic process and response to oxidative stress. The MF terms were enriched for catalytic activity and protein binding (Table S6).

**Figure 6.**
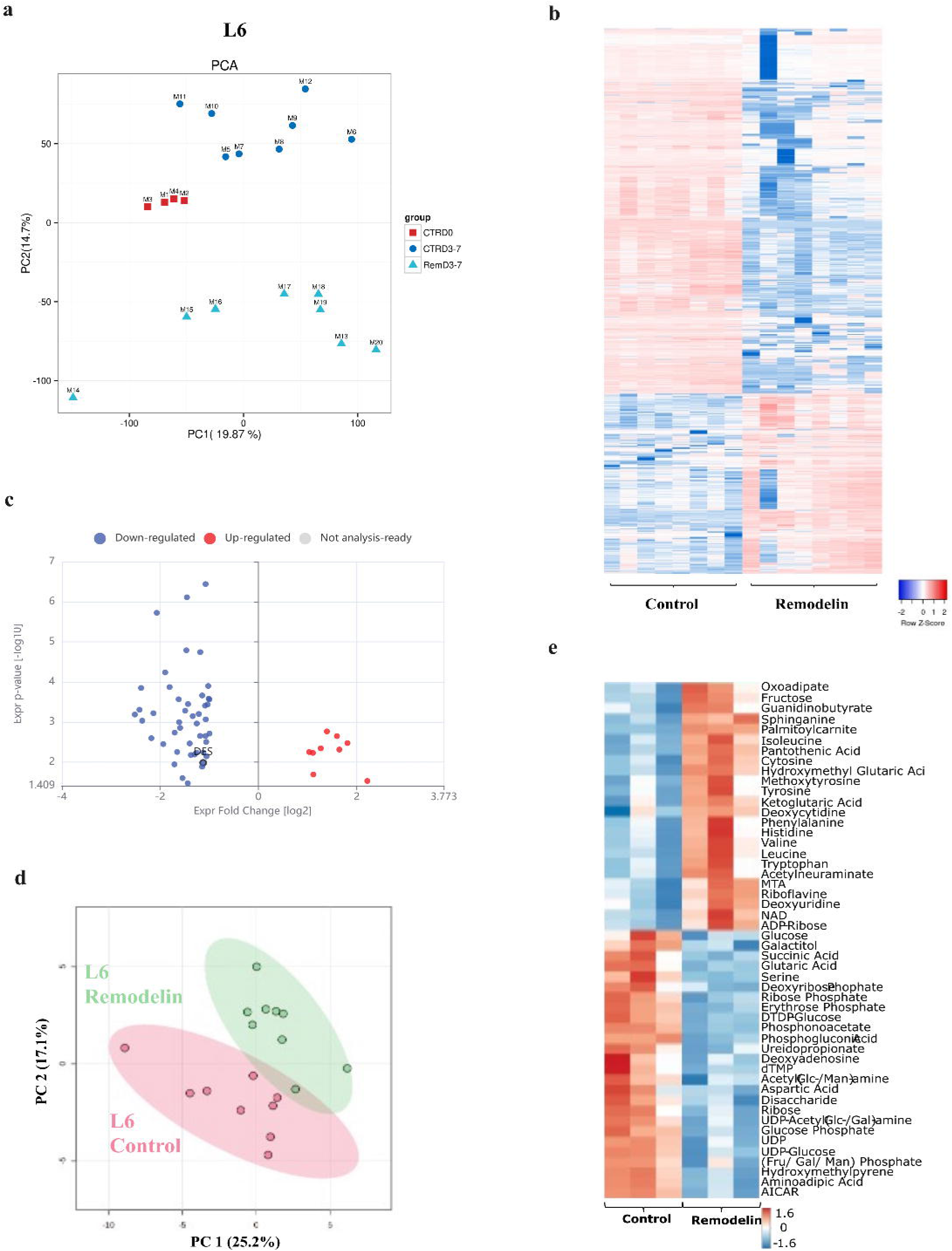
Transcriptomic, proteomic, and metabolomics profiling of L6 cells during differentiation following Remodelin treatment. (a) Principal component analysis (PCA) of RNA-seq data from L6 cells at day 0, day 3, and day 7 of differentiation, showing separation of control and Remodelin-treated samples along PC1 (19.87%) and PC2 (14.7%), indicative of treatment-dependent transcriptional reprogramming. (b) Heatmap of differentially expressed genes in control and Remodelin-treated cells at day 3-7. In blue the down-regulated genes, and in red the up-regulated genes. (c) Volcano plot of differentially expressed proteins (DEPs) at day 7 of differentiation representing the up-regulated (in red) and down-regulated (in blue) proteins. (d) Principal Component Analysis of metabolomics analysis at day 7 of differentiation, depicting clusterization of control and treatment groups along the PC1 (25.2%) and PC2 (17.1%). (e) Heatmap of significantly altered metabolites in control and Remodelin-treated groups.

Proteomics analysis was performed on L6 cells at day 7 of differentiation with and without Remodelin treatment. The analysis identified 10 up-regulated and 47 down-regulated terms in the Remodelin samples as shown in the volcano plot (Fig. 6c). gProfiler analysis revealed that downregulated proteins were mostly enriched for GO terms associated to the extracellular matrix, sarcoplasmic reticulum, myofibril (Fig. S7a). In contrast, the upregulated proteins were enriched for GO terms related to the lipid and glucose transport (Fig. S7b).

Metabolomic profiling was performed at day 7 of differentiation. PCA showed partial overlap between control and Remodelin-treated samples, separated along the PC1 (25.2%) and PC2 (17.1%) (Fig. 6d). The partial overlap reflects the relatively mild effects induced by Remodelin on L6 differentiation. The heatmap of the 50 most significant altered metabolites showed the different expression in the two groups (Fig 6e). Analysis of selected metabolites, including glucose, uric acid, NADPH, UDP-glucuronic acid, acetylcarnitine, and lactate, is presented as boxplots (Fig. S8). Interestingly, the expression trends of these metabolites in L6 cells were opposite to those observed in C2C12 cells, highlighting model-specific differences in metabolic response to Remodelin.

### Remodelin effects on histones and protein markers of differentiation in L6 cells

Western blot analyses for histone markers at day 7 of differentiation revealed that Remodelin treatment induced a down-regulation of histones H3K9/14ac (p<0.01) and H3K27me2 (p<0.01), while the other histone markers of acetylation and methylation did not change compared to the control (Figure 7a-b).

**Figure 7.**
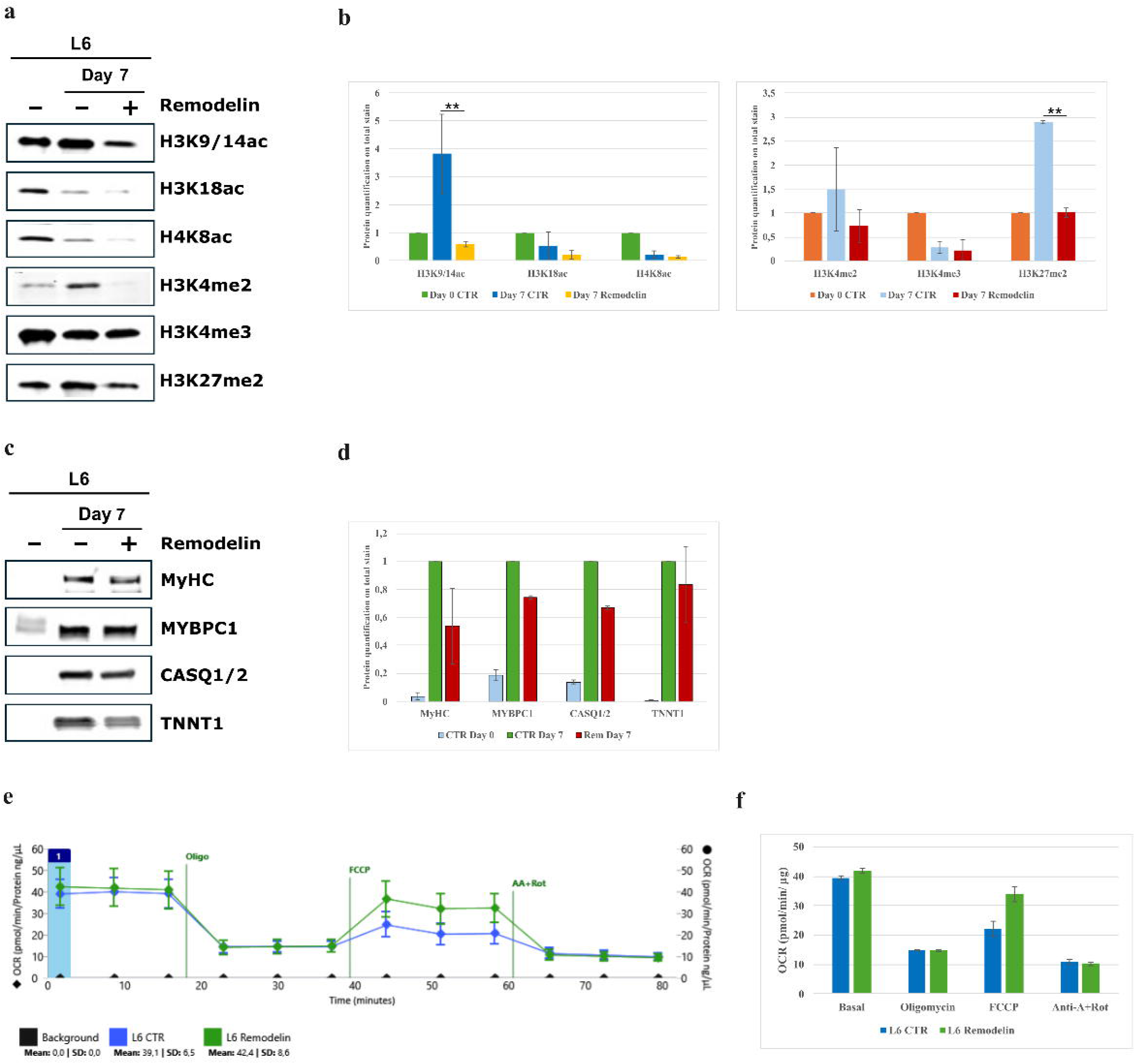
Western blot analyses of histone marks and markers of differentiation in L6 cells. (a) Representative western blots for histone acetylation marks (H3K9/14ac, H3K18ac, H4K8ac) and histone methylation marks (H3K4me2, H3K4me3, H3K27me2) in L6 cells at day 7 of differentiation, in control and Remodelin-treated cells. (b) Histograms representing histone quantification normalized on total protein stain. Histone levels are expressed relative to control cells at day 0. Data are presented as mean ± SEM (n = 3). Statistical significance is indicated as follows: ** p ≤ 0.01. (c) Representative western blots of differentiation markers at day 7 of differentiation — Myosin heavy chain II (MyHC), Mybpc1, Calsequestrin 1/2 (CASQ1/2), and Troponin T (TNNT1) — in control and Remodelin-treated cells. (d) Bar chart representing the normalization of proteins on total protein stain. Control day 7 was considered as reference control. Data are representative of three independent experiments. (e) Measurements of oxygen consumption rate (OCR) before (basal) and after addition of oligomycin, FCCP, and antimycin A in combination with rotenone, in L6 myoblasts after 24 h of Remodelin 25 μM treatment. (F) Averaged OCR values of two biological experiments. Data are presented as mean±SD (n=30 for control and n=30 for Remodelin samples).

Western blot analysis of key differentiation markers (MyHC, MYBPC1, CASQ1/2, and TNNT1) revealed a slight reduction in their expression following Remodelin treatment at day 7 of differentiation. This trend closely mirrors the effects observed in C2C12 cells and further supports the conclusion that Remodelin impairs the myogenic differentiation program (Fig 7c-d).

### Mitostress test in L6 myoblasts

We performed the mitostress test in L6 myoblasts to assess the effects of Remodelin on mitochondrial respiration. We treated the cells for 24 h with Remodelin 25 µM prior the assay. After sequential addition of oligomycin, FCCP, and antimycin A in combination with rotenone, OCR parameters were assessed. Remodelin treatment did not significantly alter basal respiration, proton leak, or non-mitochondrial respiration in L6 cells after 24 h of treatment (Fig 7e-f).

## Discussion

To the best of our knowledge, this represents the first study to investigate the impact of an epi-drug on skeletal muscle differentiation by cultivating C2C12 myoblasts on a 2D scaffold system up to 16 days. Cultivating and differentiating skeletal muscle cells has been challenging due to myotube delamination from synthetic substrate after only a few days in culture (22). The ultra-compliant gelatin hydrogel designed by Jensen and colleagues (23), together with the differentiation protocol recently optimized by our group (24), overcame this limitation in the present study by enabling sustained differentiation for at least 16 days and supporting the formation of fully mature, contractile myotubes in untreated conditions.

To assess the effects of Remodelin, a small molecule inhibitor of Nat10, on myogenic differentiation, we compared two myoblasts cell lines, C2C12 murine myoblasts and L6 rat myoblasts. L6 cells differ from C2C12 cells in terms of substrate adherence, differentiation kinetics, transcriptomic and metabolic profiling (25). Our findings demonstrate that, in both C2C12 and L6 cells, Remodelin treatment impaired myogenic differentiation, although with attenuated effects on L6 cells. In C2C12 cells, Remodelin caused a pronounced delay in myoblast fusion, severe defects in myotube alignment, and persistent structural disorganization throughout long-term differentiation. These morphological abnormalities were accompanied by a complete loss of spontaneous myotube twitching, indicating a failure to acquire functional contractile properties. In contrast, L6 myoblasts exhibited milder differentiation impairment, characterized by altered myotube morphology and disrupted nuclear organization without complete abrogation of differentiation markers at day 7 of differentiation. These observations highlight a shared sensitivity to Remodelin during myogenesis, while underscoring intrinsic differences in differentiation capacity between the two cell lines. Immunofluorescence analyses showed that nuclear positioning was disrupted in both models upon Remodelin treatment, with increased nuclear clustering and loss of the characteristic longitudinal nuclear distribution observed in control myotubes. The effect of this change in nuclear distribution may affect protein turnover, resulting in decreased production of contractile proteins and consequently compromised muscle functionality (26,27).

Multi-omics analyses provided mechanistic insight into these phenotypic alterations. Remodelin induced convergent transcriptional and proteomic alterations in both cell lines, particularly affecting pathways related to sarcomere organization, cytoskeletal dynamics, extracellular matrix (ECM) interactions, and calcium handling, processes crucial for proper muscle cell maturation and differentiation. In C2C12 cells, these changes were extensive and sustained, encompassing downregulation of genes and proteins involved in muscle contraction, excitation–contraction coupling, and ECM organization. L6 cells showed similar pathways affected, including cytoskeleton, myofibril, and ECM-related genes and proteins, but with smaller fold changes, fewer deregulated targets, and delayed onset. However, it cannot be excluded that the lower number of deregulated proteins observed in L6 cells reflects, at least in part, a technical bias due to differences in proteome annotation depth between rat and mouse databases used for mass spectrometry analysis. In C2C12 the downregulation of genes and proteins involved cytosolic calcium is crucial for both inducing myogenic differentiation and contraction, and recent findings suggest its role in myoblast-to-myotube fusion (28). Calcium plays a pivotal role in myogenesis, and its observed deregulation may well justify the differentiation slowdown and the impaired contraction seen upon Remodelin treatment. Sarcolemma is also essential for muscle structure and function. Remodelin-dysregulated pathways also include genes involved in ECM structural constituent, collagen binding, cell adhesion, and laminin-complexes (29). ECM accomplishes several important functions in skeletal muscle, offering mechanical support for muscle development, maintenance, and contraction, as well as facilitating the activation of quiescent skeletal muscle precursor cells (30,31). Deregulation of ECM, laminin complexes, and collagen binding results in defects of skeletal muscle homeostasis and its structure, thus causing a variety of muscle-related disorders.

Metabolomic profiling of C2C12 revealed clear separation between control and Remodelin-treated samples, with alterations in glucose, NADPH, lactate, and other metabolites involved in energy metabolism and redox homeostasis. In L6 cells, metabolomics showed only modest changes, and PCA indicated partial overlap between control and treated samples Some metabolites, including Acetylcarnitine shown in Fig. S5, exhibited trends opposite to those in C2C12 highlighting cell-line specific metabolic responses. Seahorse mitostress analyses confirmed that short-term Remodelin treatment did not significantly affect basal or maximal OCR in either cell line.

In support, protein levels of markers of differentiation, such as Mybpc1, CASQ1/2, myosin heavy chain, and troponin T1 were strongly impaired upon Remodelin treatment in C2C12 but only modestly decreased in L6 cells. Their combined downregulations confirm the negative effect of Remodelin in myotube formation and contraction.

We also observed a reduction in histone acetylation caused by drug treatment during the differentiation, suggesting that the effects of Remodelin might be due to the inhibition of Nat10 acetyltransferase activity. The three acetylated histones analyzed in this study – H3K9/14ac, H3K18ac, H4K8ac – are associated with transcriptional activation. Consequently, the decrease in the acetylation levels may have caused the transcriptional inactivation of genes involved in muscle differentiation.

Notably, Nat10 expression exhibited a dynamic pattern during differentiation in both cell lines, with high levels in proliferating myoblasts, a decline during early and intermediate differentiation, and a subsequent increase at late stages corresponding to myotube maturation in C2C12 cells. This conserved expression profile suggests that Nat10 activity may be tightly regulated during myogenesis and required at specific stages of muscle development.

Collectively, our findings highlight Remodelin as a potent modulator of myotube maturation and functional differentiation, with Nat10 emerging as a potential epigenetic regulator of skeletal muscle development. These observations also indicate that the biological outcome of Nat10 inhibition is highly context- and stage-dependent, suggesting that while epigenetic modulation may support muscle protection under stress conditions (19), it may conversely interfere with the chromatin remodeling programs required for effective myogenic differentiation. More broadly, this study suggests the importance of epigenetic mechanisms in muscle differentiation and suggests that epidrugs may represent a promising, yet complex, therapeutic avenue for muscle-related disorders. In the context of sarcopenia and other muscle-wasting conditions, where impaired differentiation, reduced regenerative capacity, and epigenetic dysregulation have been reported, targeting epigenetic regulators may offer new opportunities to modulate muscle plasticity.

## Acknowledgements

We thank the HiLife Flow Cytometry Unit, University of Helsinki where flow cytometry analysis was performed. We also thank Saara Tegelberg for the assistance with immunofluorescence. We thank BMKGene (Biomarker Technologies) for RNAseq. VS received funding from the “XXXVI ciclo - Dottorato di ricerca in scienze biochimiche e biotecnologiche”. AR received funding from the Deutsche Gesellschaft für Muskelkranke (DGM). AH acknowledges the support by the “Ministerium für Kultur und Wissenschaft des Landes Nordrhein-Westfalen” and “Der Regierende Bürgermeister von Berlin, Senatskanzlei Wissenschaft und Forschung” and the Bundesministerium für Forschung, Technologie und Raumfahrt (BMFTR). AN acknowledges the support by PRIN2020 project, “2020CW39SJ, Mechanistic understanding of NADPH oxidases and their roles in ROS biology” and PNRR-MAD-2022-12376672 project, “Dissecting the molecular and cellular pathophysiology of sarcopenic obesity in the elderly”. KWS acknowledges funding from Federal Ministry of Research, Technology and Space (Bundesministerium für Forschung, Technologie und Raumfahrt; BMFTR) under the funding reference 161L0271 and support by the Ministry of Culture and Science of the State of North Rhine-Westphalia (Ministerium für Kultur and Wissenschaft des Landes Nordrhein-Westfalen, MKW NRW). PEG and KWS also acknowledge the support of the EFRE/JTF program 2021-2027 (B2B-RARE; EFRE-20800340). MS, BU acknowledge the support by Samfundet Folkhälsan i Svenska Finland, The Research Council of Finland, and Sigrid Jusélius foundation. PHJ acknowledges Medicinska Understödsföreningen Liv och Hälsa rf. During the preparation of this work the authors used ChatGPT in order to proofread the final draft. After using this tool/service, the authors reviewed and edited the content as needed and take full responsibility for the content of the publication.

## Author contributions

Conceptualization: VS, MS, AN

Data curation: VS, MS, AN, AR,

Formal Analysis: VS, AH, PEG, KWS

Funding acquisition: AR, AH, BU, MS, AN

Investigation: VS, AH, PEG, JS, PHJ, KWS, SNG

Resources: AM, DR

Supervision: JS, PHJ, OE, BU, LA, MS, AN

Writing – original draft: VS

Writing – review & editing: ALL

## Data availability

RNAseq BAM files have been uploaded in NCBI SRA (accession ID: PRJNA1336407, PRJNA1423193).

‘LC-MS metabolomics raw data will be publicly available upon publication at the MassIVE repository: MassIVE MSV000100836. Reviewer access can be requested during manuscript review.

## Material and methods

### Cell culture

C2C12 murine myoblasts were purchased from ATCC (CRL-1772). Cells were mycoplasma free. Cells were cultured in Dulbecco’s modified Eagle medium (DMEM, Gibco), without phenol-red and pyruvate, supplemented with 20% fetal bovine serum (SERANA S-FBS-SA-015), 1× glutamax, and 1× penicillin/streptomycin in incubator at 37°C under humidified atmosphere of 5% CO_2_. Cells were subcultured when they reached 50% of confluence.

Ultra-compliant gelatin hydrogels were prepared essentially as described (23,24). For the differentiation, C2C12 myoblasts were grown on gelatin hydrogels to confluency and differentiated to myotubes up to 16 days in differentiation medium (DMO) containing pyruvate-free DMEM with 2% heat-inactivated horse serum, 1× L-glutamine, 1× penicillin/streptomycin and 10% Opti-MEM (24). DMO was changed every day. Collagenase type I 1 mg/mL (Gibco) was used to collect samples at day 3, 7, and 16 of differentiation.

L6 rat myoblasts (ATCC-CRL-1458) were purchased from LGC Nordic. Cells were mycoplasma free. Cells were cultured in Dulbecco’s modified Eagle medium (DMEM, Gibco), without phenol-red and pyruvate, supplemented with 10% fetal bovine serum, 1× glutamax and 1× penicillin/streptomycin in incubator at 37°C under humidified atmosphere of 5% CO_2_. Cells were subcultured when they reached 50% of confluence.

For differentiation, L6 myoblasts were grown on collagen coated (Collagen bovine, type I, Corning) plates. DMO media was changed every day. Samples were collected after 3 and 7 days of differentiation.

### Drug

Remodelin, synthesized as reported in Ref 15, was dissolved in dimethyl sulfoxide (DMSO). The drug was used at a final concentration of 25 µM or as stated. Cells were treated every 48 h and media was changed every 24 h.

### Cytotoxicity assay

The 3-(4,5-dimethylthiazol-2-yl)-2,5-diphenyl-2H-tetrazolium bromide (MTT) assay was performed to evaluate the C2C12 and L6 cell viability upon Remodelin treatment. C2C12 and L6 myoblasts were treated with different concentrations of Remodelin (5, 10, 25 µM) every 48 h. MTT assay was performed at different time points after Remodelin treatment (24 and 48h), as previously described (32). Data are presented as mean±SD of three biological replicates.

### FACS analysis

FACS analysis was performed on C2C12 myoblasts after 24 h of treatments using Remodelin at 25 μM. Cells were fixed by slowly adding ice-cold 70% ethanol and kept at 4°C for 4 h. Samples were centrifuged at 500 g for 10 min and washed once with PBS + 2% FCS. Then, 100 µl RNase A 1 mg/ml was added and the samples were incubated at 37°C for 30 min. 20 μg of propidium iodide (PI) was added and samples were incubated for ≥30 min at RT protected from light. NovoCyte Quanteon 4025 Analyzer was used for cell cycle detection.

### Total protein extraction and western blotting

C2C12 and L6 cells were collected and washed three times with 1× PBS and centrifuged at 1500 g for 5 minutes at 4°C. Cell pellets were suspended in RIPA Buffer (NP-40 1%, 50 mM Tris-HCl pH 8.0, 150 mM NaCl, 10% glycerol, 1 mM EDTA) with 1× protease inhibitor cocktail (Roche). The pellets were vortexed three times every 10 minutes, kept on ice, and then centrifuged at 15000 g for 20 min at 4°C. Protein concentration was determined by Bradford assay (Bio-Rad, California, US). For western blotting analyses, protein samples were separated in TGX minigels (Bio-Rad, Hercules, CA, USA) and transferred on nitrocellulose membranes with the Trans-Blot Turbo system (Bio-Rad). Total protein was stained with Revert 520 Total Protein Stain (Li-Cor Biosciences, Lincoln, NE, USA) and scanned with Odyssey M scanner (Li-Cor). The stain was detected in the 520 nm channel for quantification and normalization analyses.

Then, the membranes were blocked with 5% milk, and 1× PBS + 0.1% Tween (PBS-T) was used for membrane washings. Primary antibodies were diluted in 1% milk and incubated at 4°C overnight. Primary antibodies used are the following: anti-Nat10 (Abcam ab251186), anti-MyHC (DSHB), anti-Mybpc1 (Abcam ab55559), anti-Calsequestrin 1/2 (Abcam ab3516), anti-Troponin T (Novocastra). After 3 washes with PBS-T for 5 minutes, membranes were incubated with fluorescent secondary antibodies (1:10000) diluted in milk 5% for 1 hour at RT. Odyssey scanner was used to detect the signals. Empiria studio (Li-Cor) was used for the analysis. All quantitative data are expressed as mean ± SD of three biological replicates. One-way Anova in GraphPad Prism 5.02 software was used for statistical analysis.

### Histone extraction and western blotting

C2C12 samples were collected 3 days after the switch to DMO, to capture early myogenic dynamics, and after 7 days and 16 days, representing an intermediate and late stage of differentiation, respectively. L6 samples were collected 7 days after the switch to DMO. Cells were collected and washed with 1× PBS, then resuspended in Triton Extraction Buffer (TEB: PBS containing 0.5% Triton X-100 (v/v), 2 mM phenylmethylsulfonyl fluoride (PMSF), 0.02% (w/v) NaN_3_) and kept on the rotator for 10 minutes. Then, samples were centrifuged at 2000 g for 10 minutes at +4°C. Supernatant was removed and the pellet washed in half the volume of TEB and centrifuged as before. Pellet was resuspended in 0.2N HCl for acid extraction over night at 4°C. Samples were centrifuged at 2000 rpm for 10 minutes at 4°C. Protein content was determined using Bradford assay. Western blot analyses were performed for three acetylation targets, namely histone 3 lysine 9/14 acetylated (H3K9/14ac, Cell signaling), histone 3 lysine 18 acetylated (H3K18ac, Diagenode), histone 4 lysine 8 acetylated (H4K8ac, Abcam), and for three methylation targets, namely histone 3 lysine 4 di-methylated (H3K4me2, Abcam), histone 3 lysine 4 tri-methylated (H3K4me3, Abcam), and histone 3 lysine 37 di-methylated (H3K27me2, Abcam) was performed. All quantitative data are expressed as mean ± SD of three biological replicates. Statistical analysis was performed as previously described.

### Transcriptomic analysis

A total of 30 C2C12 samples and 20 L6 samples were processed for transcriptome sequencing (Table S1). Total RNA was extracted with Trizol (Invitrogen). RNA concentration and purity was measured using NanoDrop 2000 (Thermo Fisher Scientific, Wilmington, DE). RNA integrity was assessed using the RNA Nano 6000 Assay Kit of the Agilent Bioanalyzer 2100 system (Agilent Technologies, CA, USA). mRNA was purified from total RNA using oligo-dT-attached magnetic beads. Sequencing libraries were generated using NEBNext UltraTM RNA Library Prep Kit for Illumina (New England Biolabs, USA) following manufacturer’s recommendations.

Library was sequenced in Illumina NovaSeq6000. Hisat2 tools software was used to map with reference genome (Mus_musculus GRCm39). About 20 million reads per sample were generated.

Differential expression analysis was performed using DESeq2 (33). P-values were adjusted using the Benjamini and Hochberg’s approach for controlling the false discovery rate. Genes with an adjusted p-value < 0.01 were assigned as differentially expressed. Gene ontology (GO) analysis of differentially expressed genes (DEGs) was performed in three categories: biological process (BP), cellular component (CC), and molecular function (MF). Sample M3 was excluded from the analysis based on data quality control. The analysis was performed using BMKCloud (www.biocloud.net) and g:GOST of g:Profiler (34).

### Gene Set Enrichment Analysis (GSEA)

GSEA analysis was performed on RNAseq data at day 7 of differentiation by using GSEA software (v. 4.3.2). The GO: BP gene sets were used as annotated gene sets. P-value < 0.05 was considered significant.

### Proteomics

A total of 4 control samples and 4 Remodelin-treated samples at day 7 of differentiation were processed for proteomics. Proteins were extracted using the lysis buffer included in the Pierce Co-Immunoprecipitation Kit (Thermo Scientific™). Protein concentration was determined by BCA assay according to the manufacturer’s protocol. Disulfide bonds were reduced by addition of 10 mM TCEP at 37 °C for 30 min, and free sulfhydryl bonds were alkylated with 15 mM IAA at room temperature (RT) in the dark for 30 min. 100 µg protein of each sample was used for proteolysis using the S-Trap protocol (Protifi) and using a protein to trypsin ratio of 20:1. Proteolysis was stopped using formic acid to acidify the sample (pH < 3.0). A total of 1 µg of each peptide sample was separated using an Ultimate 3000 Rapid Separation Liquid Chromatography (RSLC) nano system equipped with a ProFlow flow control device, coupled with a QExactive HF Orbitrap mass spectrometer (Thermo Scientific, Schwerte, Germany). Peptide concentration was performed using a trapping column (Acclaim C18 PepMap100, 100 μm, 2 cm, Thermo Fisher Scientific, Schwerte, Germany) with 0.1% trifluoroacetic acid (Sigma-Aldrich, Hamburg, Germany) at a flow rate of 20 µL/min. This was followed by reversed-phase chromatography (Acclaim C18 PepMap100, 75 µm, 50 cm) using a binary gradient (solvent A: 0.1% formic acid (Sigma-Aldrich, Hamburg, Germany); solvent B: 84% acetonitrile (Sigma-Aldrich, Hamburg, Germany) with 0.1% formic acid). The gradient was set as follows: 5% B for 3 minutes, a linear increase to 25% over 102 minutes, further increasing to 33% over 10 minutes, and a final increase to 95% over 2 minutes, followed by a decrease to 5% over 5 minutes.

For MS survey scans, data were acquired in data-dependent acquisition mode (DDA), with full MS scans ranging from 300 to 1600 m/z at a resolution of 60,000, using the polysiloxane ion at 371.10124 m/z as a lock mass. The maximum injection time was set to 120 milliseconds, and the automatic gain control (AGC) was set to 1E6. Fragmentation involved selecting the 15 most intense ions (threshold ion count above 5E3) with a normalized collision energy (nCE) of 27% per cycle, following each survey scan. Fragment ions were recorded at a resolution of 15,000 with an AGC of 5E4 and a maximum injection time of 50 milliseconds. Dynamic exclusion was set to 15 seconds.

### Proteomics Mass Spectrometry Data Analysis

All MS raw data were processed using Proteome Discoverer software version 2.5.0.400 (Thermo Scientific, Bremen, Germany) and searched in target/decoy mode against the mouse UniProt database (www.uniprot.org, downloaded on November 21, 2019) using the MASCOT and Sequest algorithms. Search parameters included precursor and fragment ion tolerances of 10 ppm and 0.5 Da for MS and MS/MS, respectively. Trypsin was specified as the enzyme, allowing up to 2 missed cleavages. Carbamidomethylation of cysteine was set as a fixed modification, while oxidation of methionine was set as a dynamic modification. Percolator was used to establish a false discovery rate (strict) of 0.01 for both peptide and protein identifications.

A Label-free Quantification (LFQ) analysis was conducted with replicates for each condition. Only proteins identified with at least 2 unique peptides were included in the analysis. Protein ratio calculation for pairwise comparisons and a t-test (background-based) with a p-value < 0.05 were used to determine significant regulation. The number of up- and down-regulated proteins were distinctly identified based on the log-fold change criteria between the two groups, and the values were set below 0.5 for the down-regulated proteins and above 2 for the up-regulated.

QIAGEN IPA Interpret (QIAGEN Inc., https://digitalinsights.qiagen.com/products-overview/discovery-insights-portfolio/analysis-and-visualization/qiagen-ipa/qiagen-ipa-interpret/) was used to generate volcano plots (35), Heatmapper was used to generate heatmap (36), STRING (https://string-db.org/) tool was used to visualize protein-protein interaction networks, g:GOST of g:Profiler (34) was used to highlight the enriched pathways of dysregulated proteins. Hierarchical clustering analysis was performed using Gene Ontology enRIchment anaLysis and visuaLizAtion (GOrilla) tool (https://cbl-gorilla.cs.technion.ac.il/) (37).

### Metabolite extraction

A total of 4 control samples and 4 Remodelin-treated samples for C2C12, and 3 controls and 3 treatment samples for L6 at day 7 of differentiation were processed for metabolomics. For Metabolite extractions, 300 µL Acetonitrile/Methanol/Water (40/40/20; *v/v/v*) mixture including 10 µL internal standards (Stable Isotope Labeled Amino Acid Mixture for Mass Spec, Merck, Burlington, USA) was added to the cell pellets. Cell lysis was done using a Bioruptor Pico (Diagenode, Liège, Belgium) at 4 °C and the sample was stored at −80 °C overnight for protein precipitation. The samples were then centrifuged (4°C, 15000 rpm, 45 min), then the metabolite containing supernatant was transferred and dried down with nitrogen. The protein pellets were resolved in 100 µL SDS Lysis Buffer (containing 1% PhosStop, 1% Protease Inhibitor, 5% SDS) and the protein concentration was determined using a commercially available BCA-Kit (Pierce BCA Protein Assay Kit, Thermo Scientific, Waltham, MA, USA). Metabolites were reconstituted in 100 µL Acetonitrile/Water (70/30, *v/v*), vortexed (4°C, 1400 rpm, 20 min), incubated at −80 °C for 2 h and centrifuged (4°C, 15000 rpm, 60 min). The supernatant was taken and analyzed using LC-MS/MS.

### Metabolomic analysis

Metabolomic analysis was performed with an Vanquish Duo UHPLC-System coupled to an Orbitrap Eclipse Mass Spectrometer (Thermo Scientific, Waltham, MA, USA). The LC-System was operated with an Atlantis Premier BEH Z-HILIC column (5 μm, 95 Å, 100 × 2.1 mm) and the following parameters: 3 µL injection volume; 40 °C column temperature; mobile phase A: Water containing 10 mM ammonium acetate and 0.1 % ammonium hydroxide (pH 8.5); mobile phase B: Acetonitrile; 400 µL/min flow rate. The gradient parameters were: 95 % B at 0–2 min; 95–80 % B at 2–7.7 min; 80–70 % B at 7.7–9.5 min; 70–10 % B at 9.5–12 min; 10–30 % B at 12–16 min; 30–95% B at 16.5–19 min. The HESI-source was operated with the following parameters: 3500 V (positive)/ 3000 V (negative) Spray Voltage; 50 Arb Sheat Gas; 15 Arb Aux Gas; 1 Arb Sweep Gas; 330 °C Ion Transfer Tube Temperature; 350 °C Vaporizer Temperature. MS-Data were acquired in data dependent mode. The Orbitrap was operated with a scan range of 60-850 *m/z*, a resolution of 120,000 in MS1 and a resolution of 15,000 in MS2. Fragmentation was done using high energy collision dissociation (HCD) with 10, 35 and 50 % HCD collision energies.

### Metabolomic Data Analysis

Progenesis QI v3.1 (Waters Corporation, Milford, USA) was used for data analysis. Data alignment was done using pooled samples and normalization was performed using internal standards (acceptance threshold at 30% coefficient of variation). Metabolite identification was done according to isotopic masses and their retention time, which was matched with an in-house library (38). For MS2 fragment spectra the METLIN MS/MS Library 2017 was used.

Alignment of metabolite adducts was checked, with incorrect alignments corrected and summed manually. For metabolites that were identified in both positive and negative polarity, reported values for one polarity were chosen according to their Fragmentation Score and Abundance. Only Peaks detected in at least 50% of C2C12 or L6 were accepted for the respective cell type.

Statistical analysis was performed using Metaboanalyst 6.0 (https://www.metaboanalyst.ca/). Sample normalization was done using total protein amount determined by BCA. Metabolite data was Log(10) transformed and scaled by Pareto Scaling.

### Cellular mitochondrial stress

Metabolic status was investigated on a Seahorse Pro XF Analyzer (Agilent Technologies, Santa Clara, CA, USA). Mito Stress Test Kit (Agilent Technologies, #103015) was used according to Agilent protocol. Briefly, 4×10^3^ C2C12 and 8 ×10^3^ L6 cells per well were plated 48 h prior to analysis, and treated for 24 h with Remodelin 25 µM. The steps for incubation, medium replacement and loading into XF96 Analyzer and injection were performed according to protocol. The analyzer injected the following compounds: for C2C12 Oligomycin 1,5 µM, FCCP 2 µM, Antimycin A and Rotenone 1 µM were used for the assay. For L6 Oligomycin 1,5 µM, FCCP 3 µM, Antimycin A and Rotenone 0,5 µM were used for the assay. OCR and ECAR were measured. Data were normalized on the protein concentration. Data were analyzed with Wave Pro software (Seahorse Bioscience, Agilent Technologies). Values are reported as average of two biological experiments. Standard deviations reported as error bars.

### Immunofluorescence and differentiation index (DI)

C2C12 and L6 cells were permeabilized with 0.02% Triton X-100 for 10 min at RT, and washed three times with 1× PBS. Samples were blocked with BSA 5% for 1 h at RT, and then incubated with primary antibodies overnight at 4°C. Secondary antibodies were incubated for 1 h at RT protected from light. Hoechst was added for 5 min at RT. C2C12 cells on hydrogel were mounted with PVA mounting media, as previously described (24). Leica TCS SP8 STED 3X CW 3D confocal microscope and Zeiss Axio Imager M2 (Carl Zeiss AG, Oberkochen, Germany) were used for the image acquisitions.

The differentiation index (DI) is calculated in C2C12 cells at day 7 of differentiation as the percentage of nuclei located within MyHC⁺ cells versus the total number of nuclei. Quantification was performed using ImageJ software. The data are expressed as mean±SD of three independent measurements. Statistical analysis was assessed using a two-tailed unpaired Student’s t-test with Welch’s correction (unequal variance). A p-value < 0.05 was considered statistically significant.

### Statistical analysis

Data are presented as the mean ± SD as indicated in the respective material and method sections as well as figure legends. The corresponding tests and number of replicates in each experiment are indicated in the corresponding material and method sections and figure legends. A p-value below 0.05 was considered statistically significant. Statistical significance is indicated as follows: * for p < 0.05, ** for p < 0.01, *** for p < 0.001.

## Supplemental information

Supplementary Figure S1. Remodelin effects on cell viability and cell-cycle progression. (a) MTT assay evaluating C2C12 cell viability after 24, 48, and 72 h of Remodelin treatment at different concentrations (5, 10, and 25 µM). (b) Flow cytometry (FACS) analysis of cell-cycle distribution after 8 h and 24 h of Remodelin (25 µM) treatment.

Supplementary Figure S2. Effects of Remodelin in C2C12 differentiation. (a) Nat10 protein level detected in C2C12 cells after 7 and 16 days of differentiation upon Remodelin treatment (5 and 25 µM). Total protein stain was used to normalize. (b) Quantification of the Differentiation Index (DI) assessed at day 7 of differentiation in control and Remodelin-treated cells. The differentiation index (DI), calculated as the percentage of nuclei located within MyHC⁺ cells versus the total number of nuclei, was 55% in control cultures and decreased to 37% upon Remodelin treatment. The data are expressed as mean±SD of three independent measurements (p value < 0.05).

Supplementary Figure S3. Proteomic profile of dysregulated proteins in C2C12 differentiation upon Remodelin treatment. (a) gProfiler functional enrichment analysis of proteins down-regulated in response to Remodelin treatment, highlighting pathways associated with muscle structure and differentiation. (b) gProfiler functional enrichment analysis of proteins up-regulated in Remodelin-treated cells, revealing enrichment of pathways related to metabolic and stress-response processes. (c) Integrated transcriptomic and proteomic analysis showing the overlap between differentially expressed genes (DEGs) and DEPs. Venn diagrams depict commonly up-regulated and down-regulated targets shared between the two datasets.

Supplementary Figure S4. Proteomic network and functional enrichment analysis in C2C12 cells at day 7 of differentiation. (a) STRING protein-protein interaction analysis of the significantly down-regulated proteins in Remodelin group (p value < 0.05). Colored lines between the proteins indicate the various types of interaction evidence. A green line: neighborhood evidence; a blue line: cooccurrence evidence; a purple line: experimental evidence; a yellow line: text mining evidence; a black line: co-expression evidence. The analysis was performed using STRING 12.0 software (https://string-db.org/). (b) STRING analysis of significantly up-regulated proteins in Remodelin group (p value < 0.05). c) GOrilla enrichment analysis of the down-regulated proteins. The pathway analysis shows the enriched GO terms in Remodelin group, and the results are visualized using a DAG graphical representation. Colors reflect their degree of enrichment

Supplementary Figure S5. Highlighted selection of annotated metabolites detected in C2C12 cells for both control and Remodelin-treated conditions. Control and Remodelin-treated cells were compared to identify metabolite intensity changes exhibited in the box plots.

Supplementary Figure S6. Phenotypic changes in L6 differentiation following Remodelin treatment. (a) Representative phase-contrast images of L6 at day 3 and day 7 of differentiation. Scale bar 100 μm. (b) Immunofluorescence analysis of L6 at day 7 of differentiation in control and Remodelin-treated cells. In red myosin heavy chain (MyHC) and in blue the nuclei. Images were acquired at 100× magnification. Scale bar 10 μm.

Supplementary Figure S7. Proteomic analyses of L6 differentiation after Remodelin treatment. (a) gProfiler functional enrichment analysis of proteins down-regulated in Remodelin-treated L6 cells, highlighting the most significantly enriched Gene Ontology (GO) and KEGG pathways. (b) gProfiler functional enrichment analysis of proteins up-regulated upon Remodelin treatment, revealing enrichment of pathways associated with lipid and glucose transport.

Supplementary Figure S8. Highlighted selection of annotated metabolites detected in L6 cells for both control and Remodelin treated conditions. Control and Remodelin treated cells were compared to identify metabolite intensity changes exhibited in the box plots.

Video S1. Video representing twitching of control C2C12 cells at day 7 of differentiation.

Video S2. Video representing twitching of C2C12 cells treated with Remodelin (5 µM) at day 7 of differentiation.

Video S3. Video representing twitching of control C2C12 cells at day 7 of differentiation.

Video S4. Video representing twitching of C2C12 cells upon Remodelin treatment (25 µM) at day 7 of differentiation.

Video S5. Video representing twitching of control C2C12 cells at day 16 of differentiation.

Video S6. Video representing twitching of C2C12 cells upon Remodelin treatment (25 µM) at day 16 of differentiation.

Table S1. Groups of biological replicates for transcriptomic analyses in C2C12 and L6 cells.

Table S2. GO enrichment analysis of down-regulated genes at day 3 of differentiation in C2C12 cells.

Table S3. GO enrichment analysis of down-regulated genes at day 16 of differentiation in C2C12 cells.

Table S4. Relative expression levels of genes associated with muscle contraction, sarcomere organization, skeletal muscle contraction, muscle structure development, and actin filament binding in control and Remodelin-treated C2C12 cells at day 7 of differentiation.

Table S5. gProfiler enrichment analysis of Gene Ontology (GO) terms significantly down-regulated in L6 cells at day 7 of differentiation following Remodelin treatment. Enriched terms are grouped according to biological process (BP), cellular component (CC), and molecular function (MF) categories. The analysis highlights pathways associated with myogenic differentiation, cytoskeletal organization, myofibril and sarcomere assembly, extracellular matrix organization. Terms are ranked based on adjusted p-values, and only statistically significant terms are reported.

Table S6. gProfiler enrichment analysis of Gene Ontology (GO) terms significantly up-regulated in L6 cells at day 7 of differentiation following Remodelin treatment. Enriched biological process (BP), cellular component (CC), and molecular function (MF) categories reflect activation of pathways related to metabolic adaptation, ribosome biogenesis, protein synthesis, oxidative stress response, and lipid and glucose transport. Terms are ranked based on adjusted p-values, and only statistically significant terms are reported.

## Bibliography

1. Li JW, Shen ZK, Lin YS, Wang ZY, Li ML, Sun HX, et al. DNA methylation of skeletal muscle function-related secretary factors identifies FGF2 as a potential biomarker for sarcopenia. J Cachexia Sarcopenia Muscle. 2024;

2. Guimarães DSPSF, Barrios NMF, de Oliveira AG, Rizo-Roca D, Jollet M, Smith JAB, et al. Concerted regulation of skeletal muscle metabolism and contractile properties by the orphan nuclear receptor Nr2f6. J Cachexia Sarcopenia Muscle. 2024;

3. Zhu Q, Liang F, Cai S, Luo X, Duo T, Liang Z, et al. KDM4A regulates myogenesis by demethylating H3K9me3 of myogenic regulatory factors. Cell Death Dis. 2021 Jun 1;12(6).

4. Cao Y, Yao Z, Sarkar D, Lawrence M, Sanchez GJ, Parker MH, et al. Genome-wide MyoD Binding in Skeletal Muscle Cells: A Potential for Broad Cellular Reprogramming. Dev Cell. 2010 Apr;18(4):662–74.

5. Sincennes MC, Brun CE, Rudnicki MA. Concise Review: Epigenetic Regulation of Myogenesis in Health and Disease. Stem Cells Transl Med. 2016 Mar 1;5(3):282–90.

6. Bodine SC, Sinha I, Sweeney HL. Mechanisms of Skeletal Muscle Atrophy and Molecular Circuitry of Stem Cell Fate in Skeletal Muscle Regeneration and Aging. Journals of Gerontology - Series A Biological Sciences and Medical Sciences. 2023 Jun 1;78:14–8.

7. Zammit PS. Function of the myogenic regulatory factors Myf5, MyoD, Myogenin and MRF4 in skeletal muscle, satellite cells and regenerative myogenesis. Vol. 72, Seminars in Cell and Developmental Biology. Elsevier Ltd; 2017. p. 19–32.

8. Asp P, Blum R, Vethantham V, Parisi F, Micsinai M, Cheng J, et al. Genome-wide remodeling of the epigenetic landscape during myogenic differentiation. Proc Natl Acad Sci U S A. 2011 May 31;108(22).

9. Antoun E, Garratt ES, Taddei A, Burton MA, Barton SJ, Titcombe P, et al. Epigenome-wide association study of sarcopenia: findings from the Hertfordshire Sarcopenia Study (HSS). J Cachexia Sarcopenia Muscle. 2022 Feb 1;13(1):240–53.

10. Pomella S, Danielli SG, Alaggio R, Breunis WB, Hamed E, Selfe J, et al. Genomic and Epigenetic Changes Drive Aberrant Skeletal Muscle Differentiation in Rhabdomyosarcoma. Vol. 15, Cancers. MDPI; 2023.

11. Wang K, Liu H, Hu Q, Wang L, Liu J, Zheng Z, et al. Epigenetic regulation of aging: implications for interventions of aging and diseases. Vol. 7, Signal Transduction and Targeted Therapy. Springer Nature; 2022.

12. Dhar P, Moodithaya SS, Patil P. Epigenetic alterations—The silent indicator for early aging and age-associated health-risks. Vol. 5, Aging Medicine. John Wiley and Sons Inc; 2022. p. 287–93.

13. Dalhat MH, Altayb HN, Khan MI, Choudhry H. Structural insights of human N-acetyltransferase 10 and identification of its potential novel inhibitors. Sci Rep. 2021 Dec 1;11(1).

14. Rodrigues P, Bangali H, Ali E, Nauryzbaevish AS, Hjazi A, Fenjan MN, et al. The mechanistic role of NAT10 in cancer: Unraveling the enigmatic web of oncogenic signaling. Vol. 253, Pathology Research and Practice. Elsevier GmbH; 2024.

15. Zhang H, Shan W, Yang Z, Zhang Y, Wang M, Gao L, et al. NAT10 mediated mRNA acetylation modification patterns associated with colon cancer progression and microsatellite status. Epigenetics. 2023;18(1).

16. Liu HY, Liu YY, Yang F, Zhang L, Zhang FL, Hu X, et al. Acetylation of MORC2 by NAT10 regulates cell-cycle checkpoint control and resistance to DNA-damaging chemotherapy and radiotherapy in breast cancer. Nucleic Acids Res. 2021;48(7):3638–56.

17. Wei W, Zhang S, Han H, Wang X, Zheng S, Wang Z, et al. NAT10-mediated ac4C tRNA modification promotes EGFR mRNA translation and gefitinib resistance in cancer. Cell Rep. 2023 Jul 25;42(7).

18. Larrieu D, Britton S, Demir M, Rodriguez R, Jackson SP. Chemical inhibition of NAT10 corrects defects of laminopathic cells. Science (1979). 2014;344(6183):527–32.

19. Wang C, Liu Y, Zhang Y, Wang D, Xu L, Li Z, et al. Targeting NAT10 protects against sepsis-induced skeletal muscle atrophy by inhibiting ROS/NLRP3. Life Sci. 2023 Oct 1;330.

20. Shi L, Zhang M, Yang H, Li X, He S, Chu Y, et al. NAT10 regulates heart development and function by maintaining the expression of genes related to fatty acid β-oxidation and heart contraction. Cell Death Differ. 2025;

21. Shi J, Yang C, Zhang J, Zhao K, Li P, Kong C, et al. NAT10 Is Involved in Cardiac Remodeling Through ac4C-Mediated Transcriptomic Regulation. Circ Res. 2023 Dec 8;133(12):989–1002.

22. Bettadapur A, Suh GC, Geisse NA, Wang ER, Hua C, Huber HA, et al. Prolonged Culture of Aligned Skeletal Myotubes on Micromolded Gelatin Hydrogels. Sci Rep. 2016 Jun 28;6.

23. Jensen JH, Cakal SD, Li J, Pless CJ, Radeke C, Jepsen ML, et al. Large-scale spontaneous self-organization and maturation of skeletal muscle tissues on ultra-compliant gelatin hydrogel substrates. Sci Rep. 2020 Dec 1;10(1).

24. Sian V, Jonson PH, Vainio A, Luque H, Gayathri SN, Hackman P, et al. Optimizing 2D in vitro differentiation conditions for C2C12 murine myoblasts on gelatin hydrogel. J Muscle Res Cell Motil. 2025;

25. Abdelmoez AM, Puig LS, Smith JAB, Gabriel BM, Savikj M, Dollet L, et al. Comparative profiling of skeletal muscle models reveals heterogeneity of transcriptome and metabolism. Am J Physiol Cell Physiol [Internet]. 2020;318:615–26. Available from: www.ajpcell.org

26. Cisterna B, Malatesta M. Molecular and Structural Alterations of Skeletal Muscle Tissue Nuclei during Aging. Vol. 25, International Journal of Molecular Sciences. Multidisciplinary Digital Publishing Institute (MDPI); 2024.

27. Levy Y, Ross JA, Niglas M, Snetkov VA, Lynham S, Liao CY, et al. Prelamin A causes aberrant myonuclear arrangement and results in muscle fiber weakness. JCI Insight. 2018 Oct 4;3(19).

28. Sinha S, Elbaz-Alon Y, Avinoam O. Ca2+ as a coordinator of skeletal muscle differentiation, fusion and contraction. FEBS Journal. 2022 Nov 1;289(21):6531–42.

29. Gillies AR, Lieber RL. Structure and function of the skeletal muscle extracellular matrix. Vol. 44, Muscle and Nerve. 2011. p. 318–31.

30. Zhang W, Liu Y, Zhang H. Extracellular matrix: an important regulator of cell functions and skeletal muscle development. Vol. 11, Cell and Bioscience. BioMed Central Ltd; 2021.

31. Ahmad K, Shaikh S, Chun HJ, Ali S, Lim JH, Ahmad SS, et al. Extracellular matrix: the critical contributor to skeletal muscle regeneration—a comprehensive review. Vol. 43, Inflammation and Regeneration. BioMed Central Ltd; 2023.

32. Carrizzo A, Iside C, Nebbioso A, Carafa V, Damato A, Sciarretta S, et al. SIRT1 pharmacological activation rescues vascular dysfunction and prevents thrombosis in MTHFR deficiency. Cellular and Molecular Life Sciences. 2022 Aug 1;79(8).

33. Love MI, Huber W, Anders S. Moderated estimation of fold change and dispersion for RNA-seq data with DESeq2. Genome Biol. 2014 Dec 5;15(12).

34. Kolberg L, Raudvere U, Kuzmin I, Adler P, Vilo J, Peterson H. G:Profiler-interoperable web service for functional enrichment analysis and gene identifier mapping (2023 update). Nucleic Acids Res. 2023 Jul 5;51(W1):W207–12.

35. Krämer A, Green J, Pollard J, Tugendreich S. Causal analysis approaches in ingenuity pathway analysis. Bioinformatics. 2014 Feb 15;30(4):523–30.

36. Babicki S, Arndt D, Marcu A, Liang Y, Grant JR, Maciejewski A, et al. Heatmapper: web-enabled heat mapping for all. Nucleic Acids Res. 2016 Apr 1;44(1):W147–53.

37. Eden E, Navon R, Steinfeld I, Lipson D, Yakhini Z. GOrilla: A tool for discovery and visualization of enriched GO terms in ranked gene lists. BMC Bioinformatics. 2009 Feb 3;10.

38. Kasarla SS, Flocke V, Saw NMT, Fecke A, Sickmann A, Gunzer M, et al. In-vivo tracking of deuterium metabolism in mouse organs using LC-MS/MS. J Chromatogr A. 2024 Feb 22;1717.

