## Supplementary material for "The inhibitory effects of Remodelin on myoblasts differentiation": Supplemnetary Figures

**Supplementary Figures**

**Supplementary Figure S1**

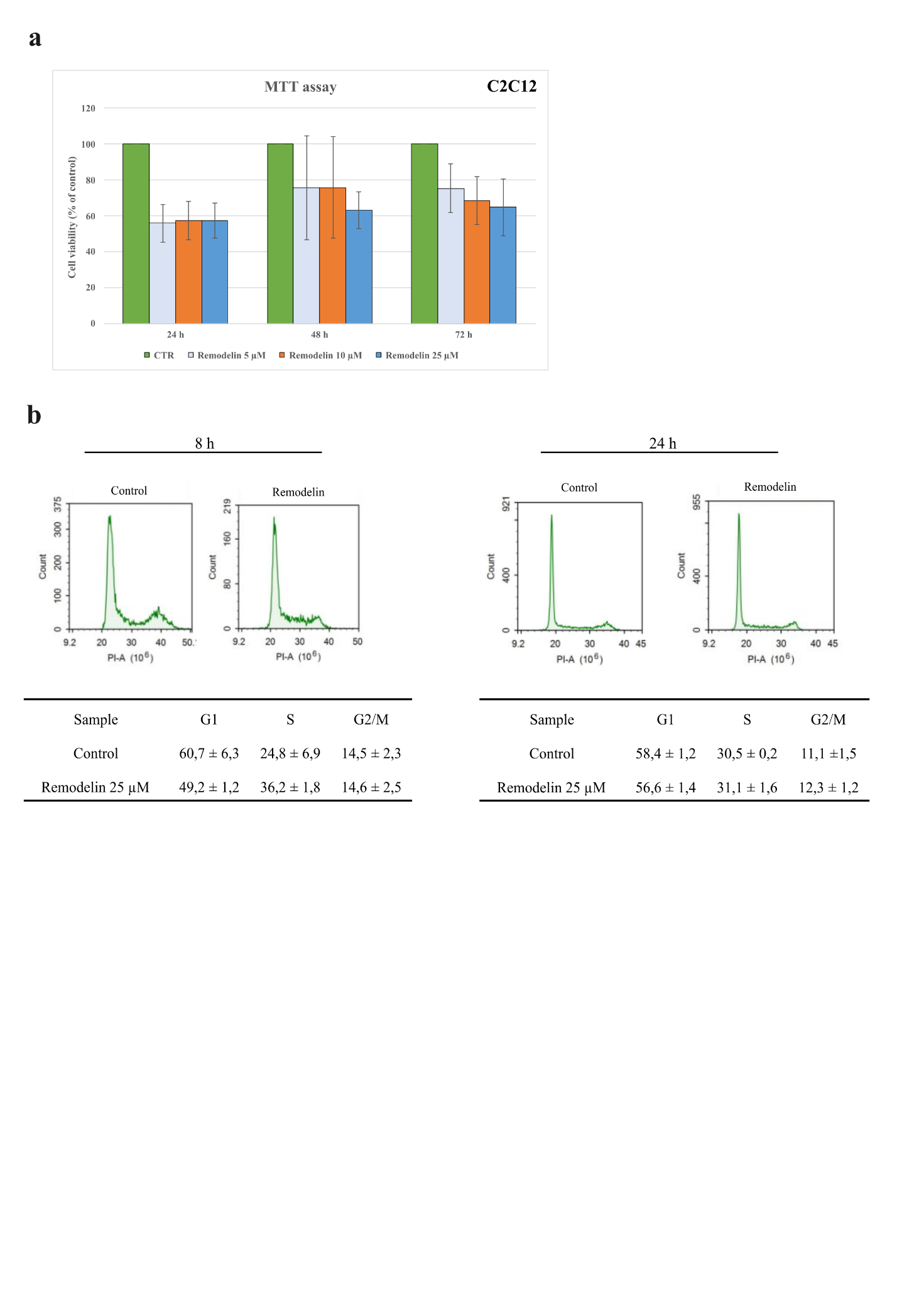

**Supplementary Figure S2**

**a**

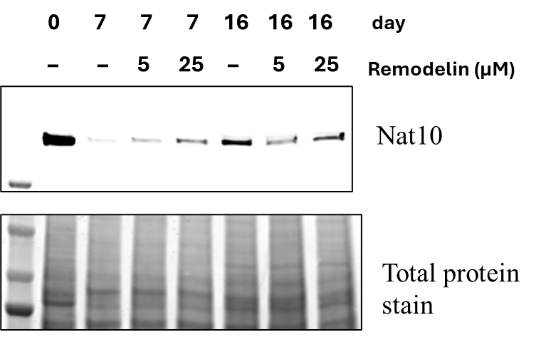

**b**

Differentiation index

% Differentiation index

|  | DI % |
| --- | --- |
| CTR | 55.0733 |
| Remodelin | 33.1033 |

**Supplementary Figure S3**

**a**

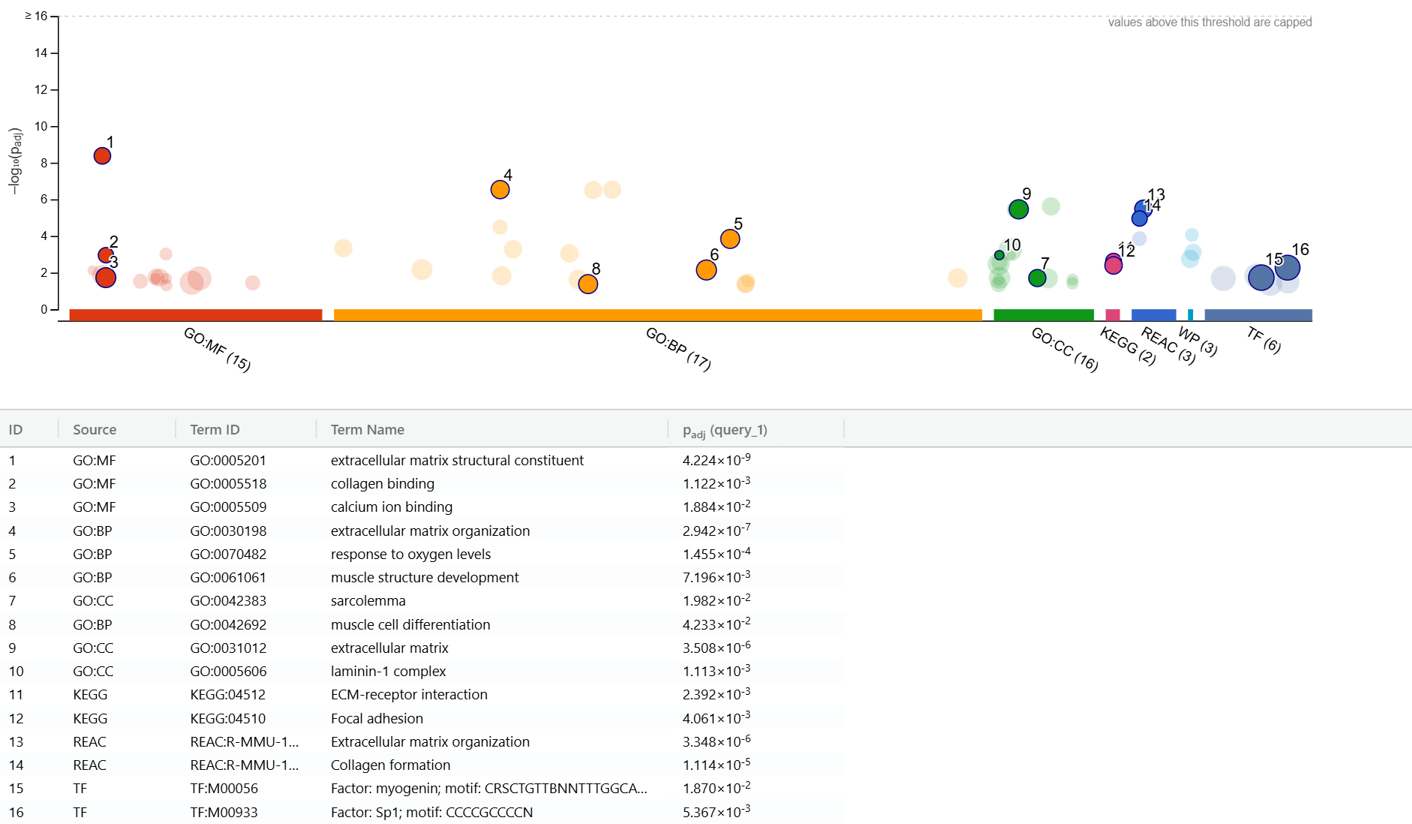

**b**

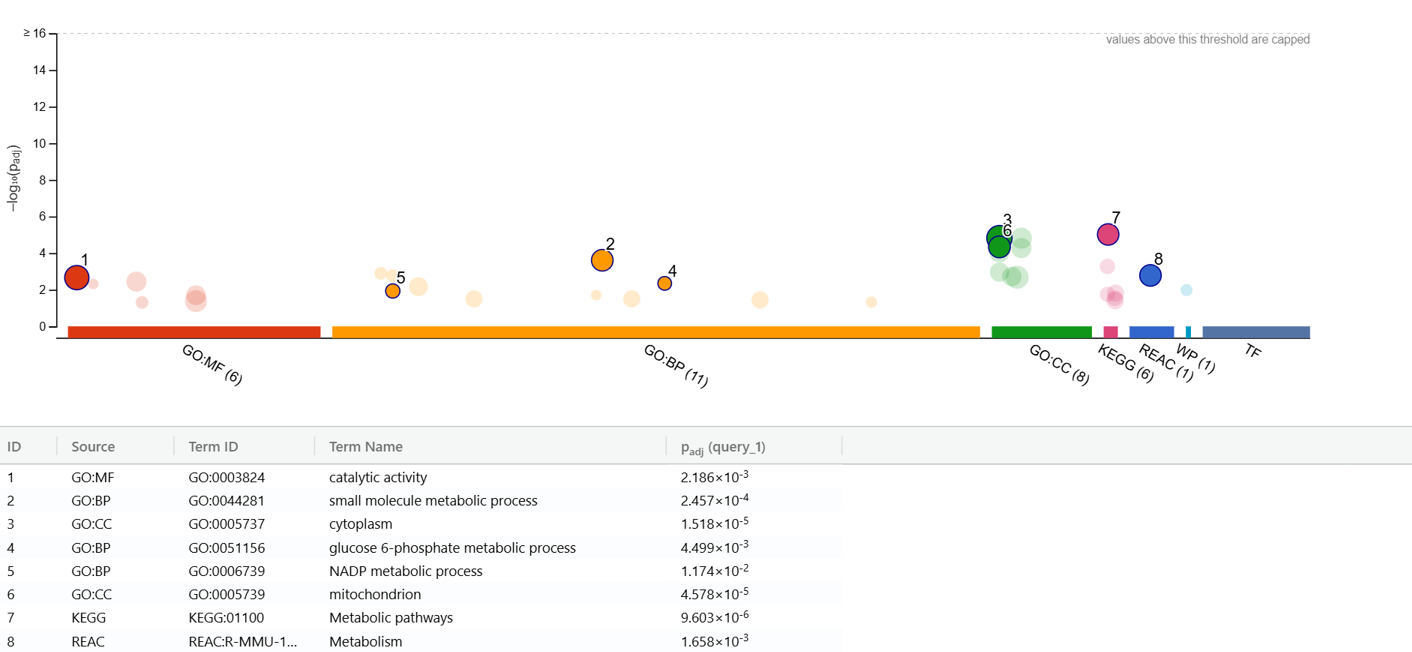

**c**

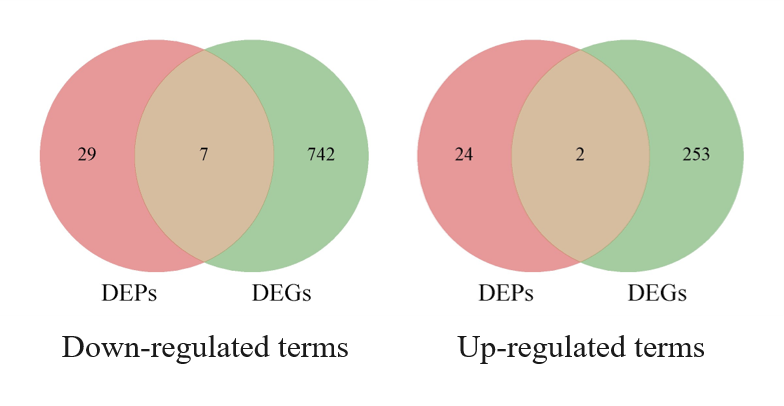

**Supplementary Figure S4**

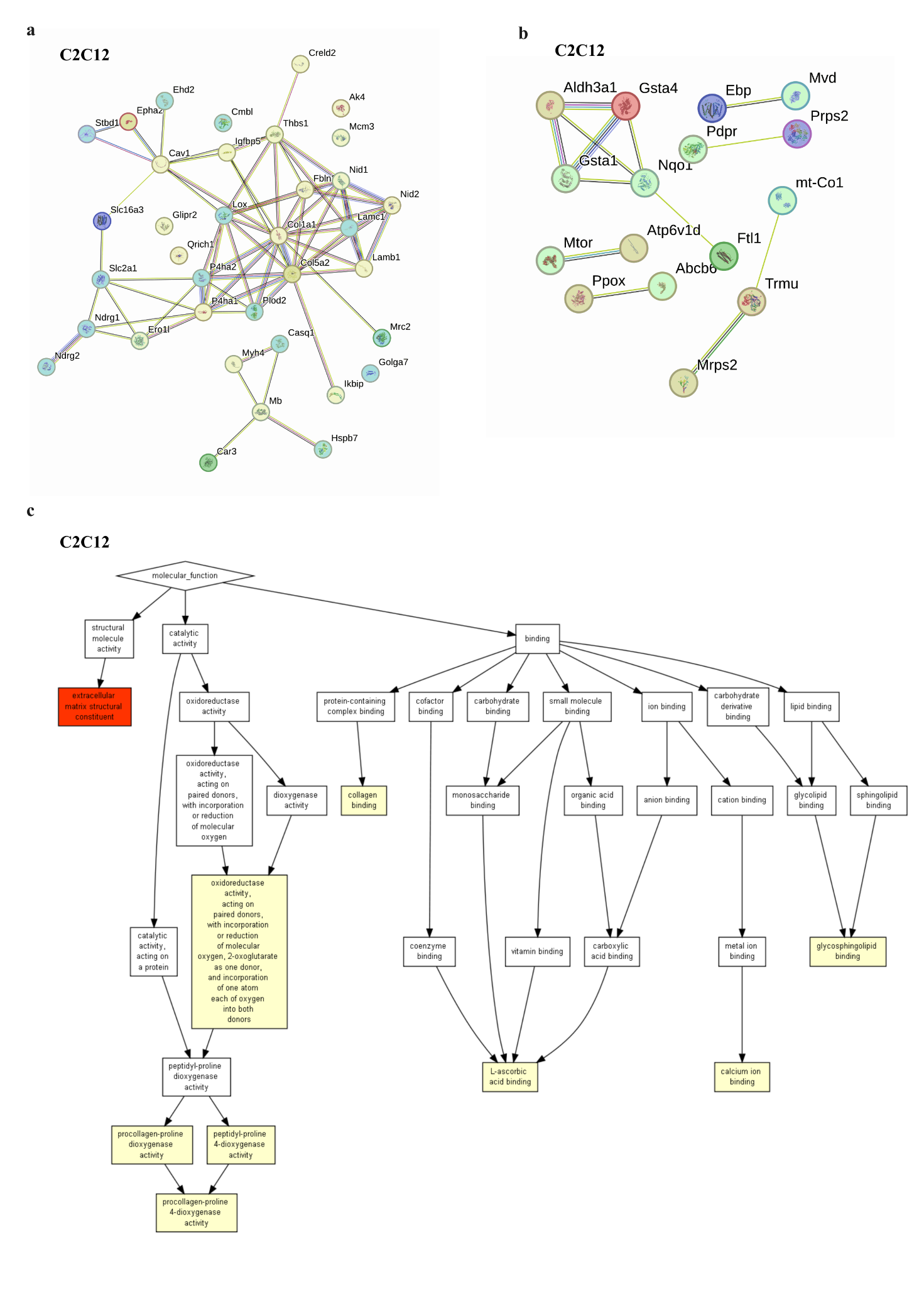

**Supplementary Figure S5**

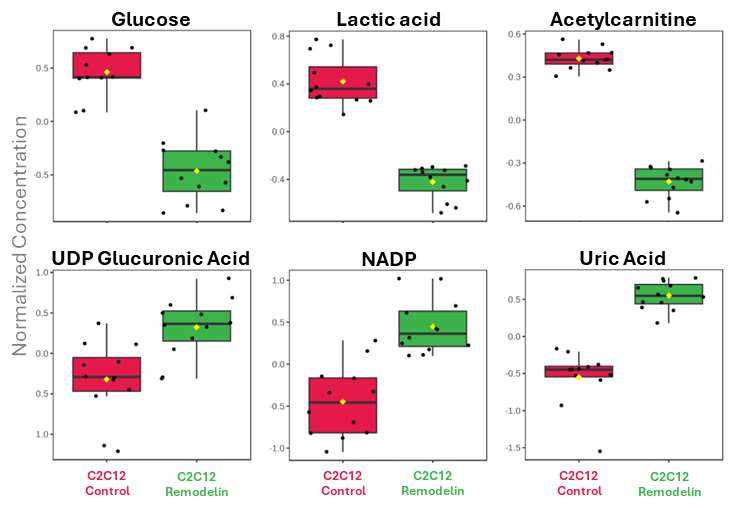

**Supplementary Figure S6**

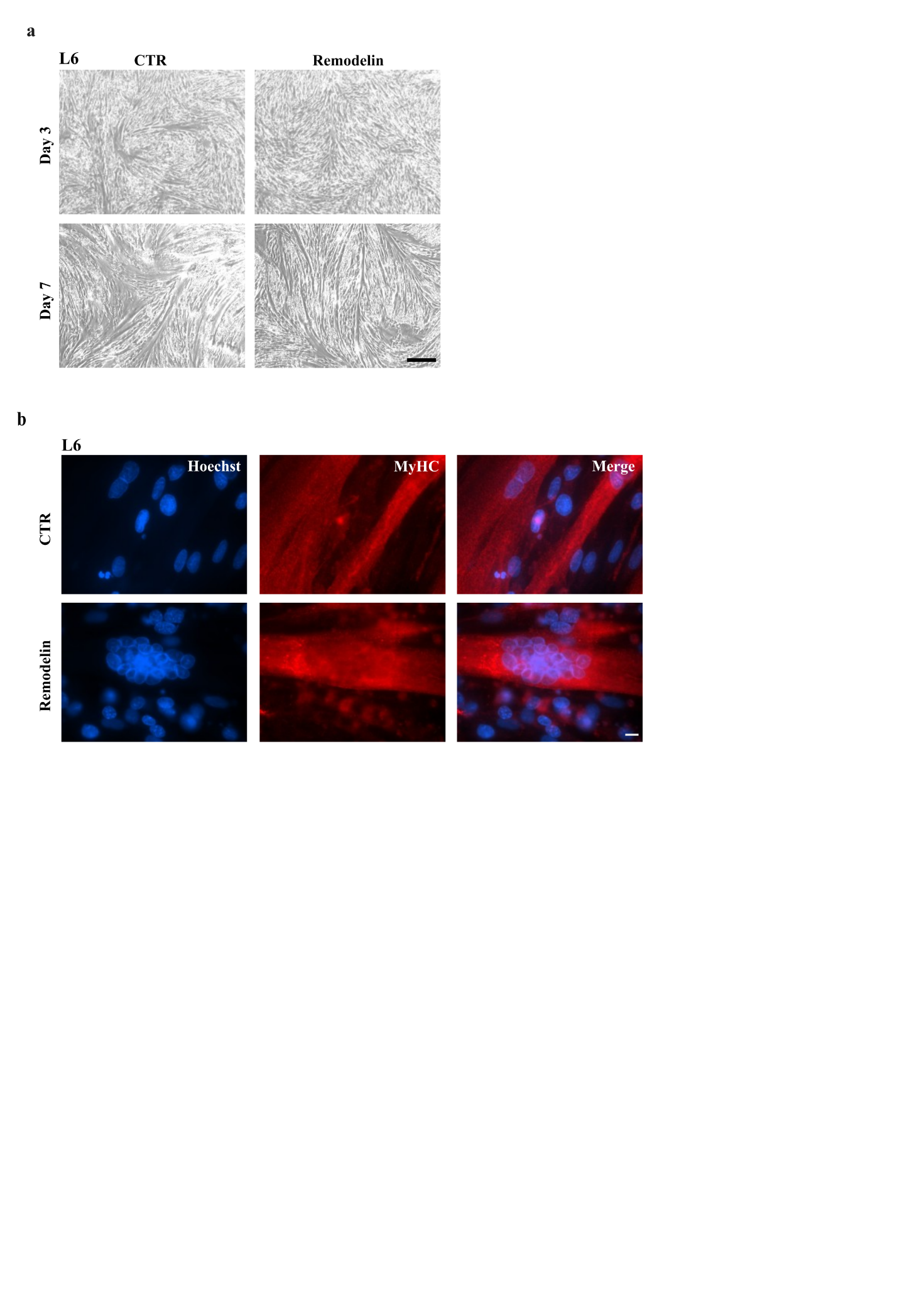

**Supplementary Figure S7**

**a**

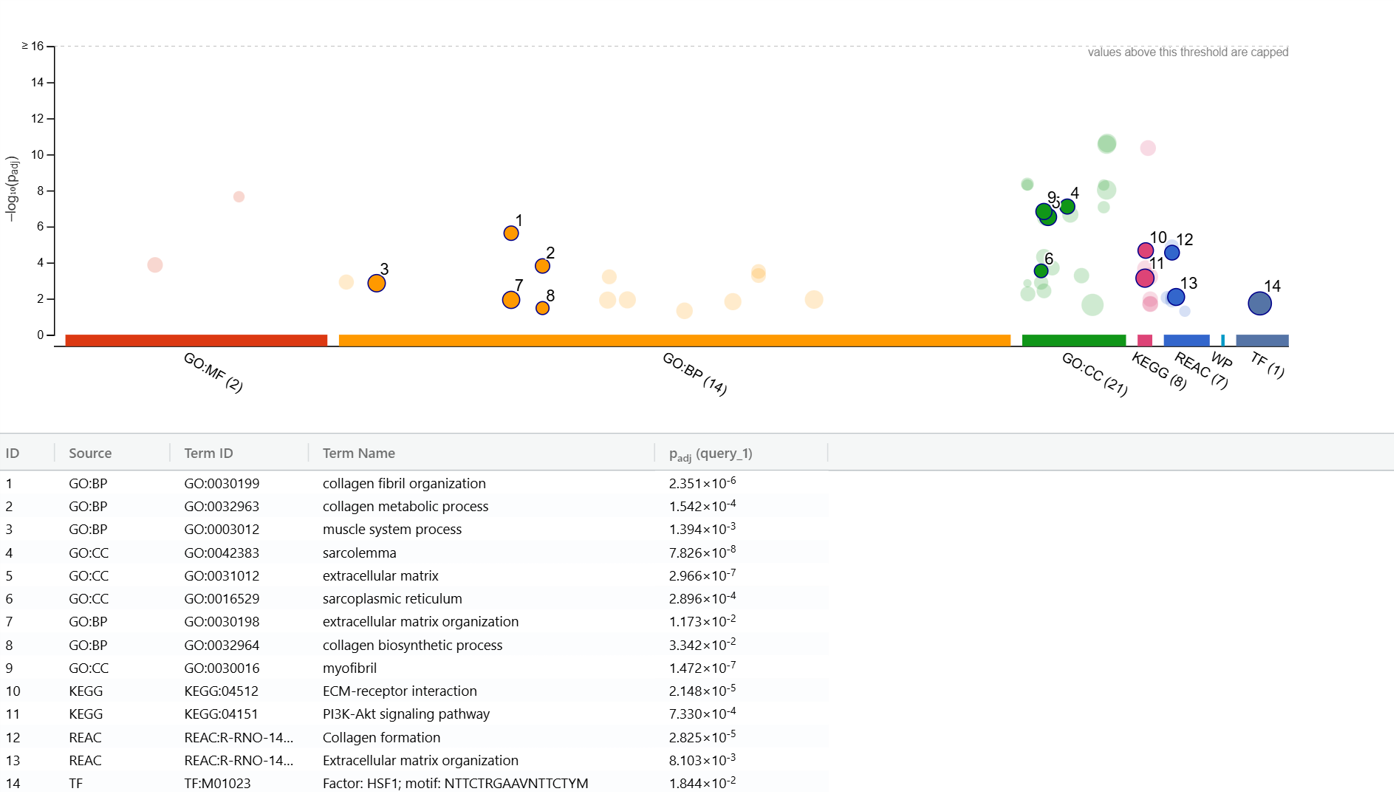

**b**

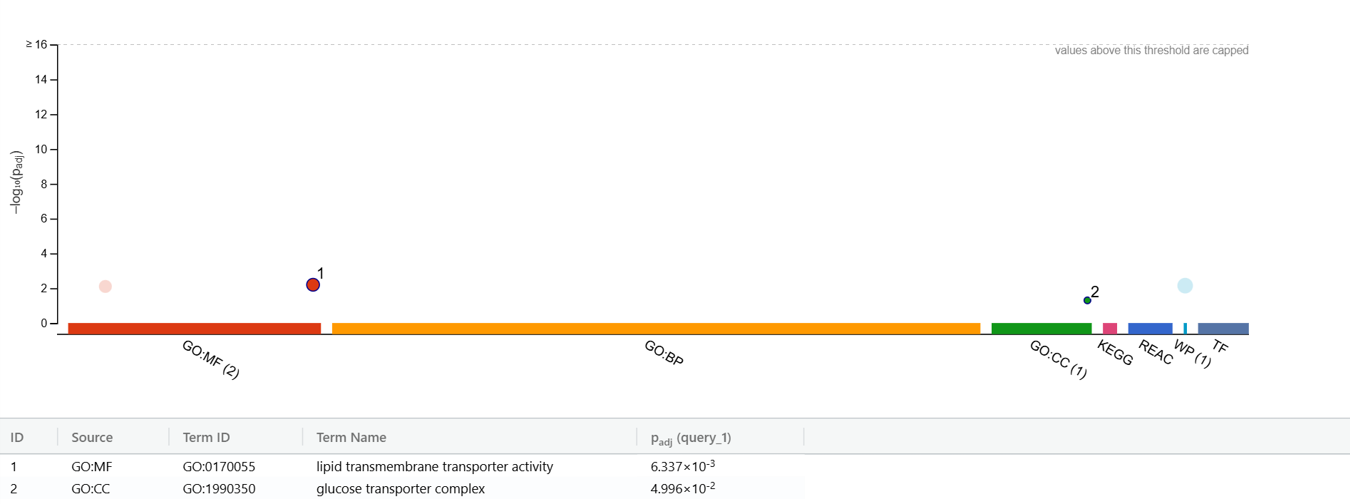

**Supplementary Figure S8**

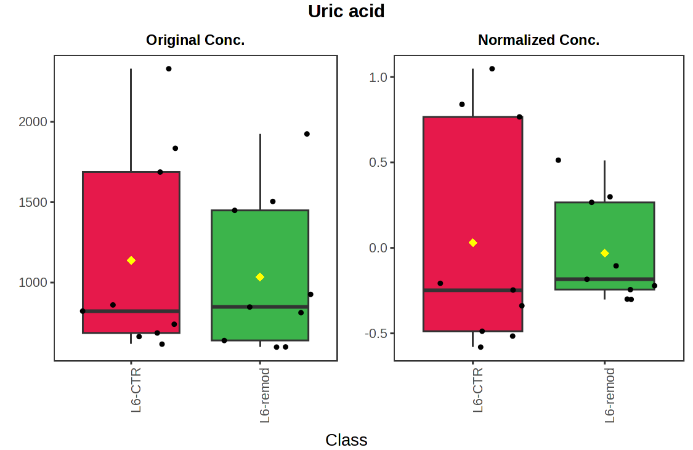

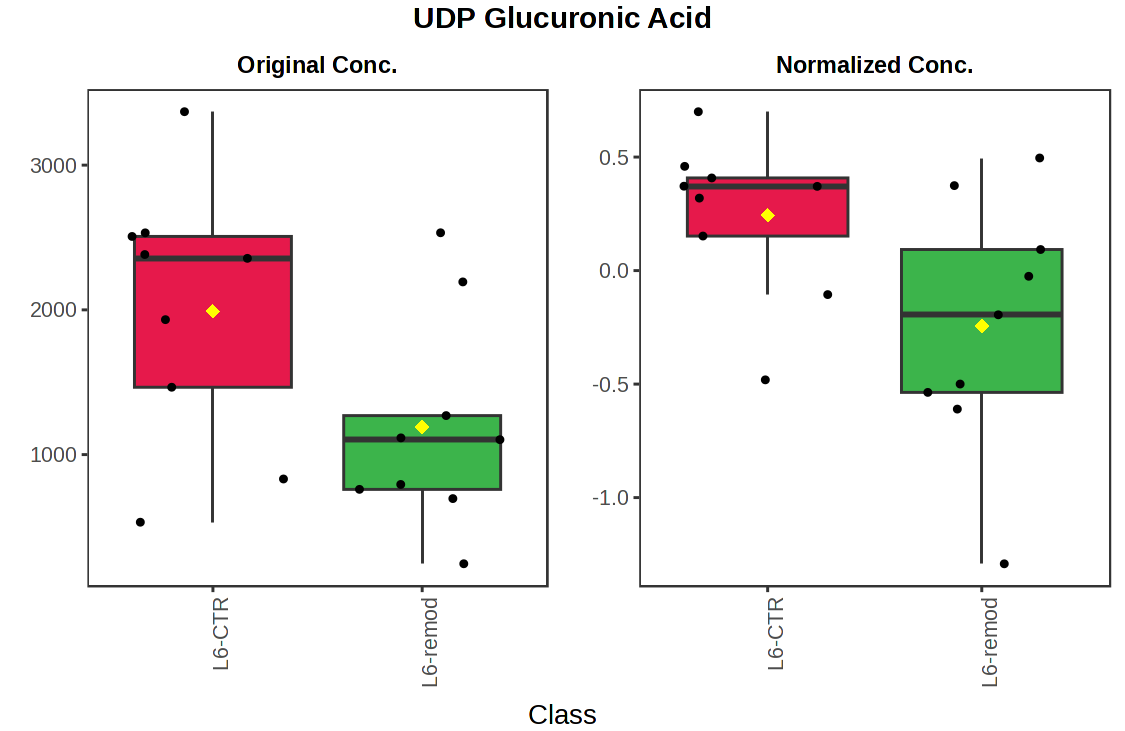

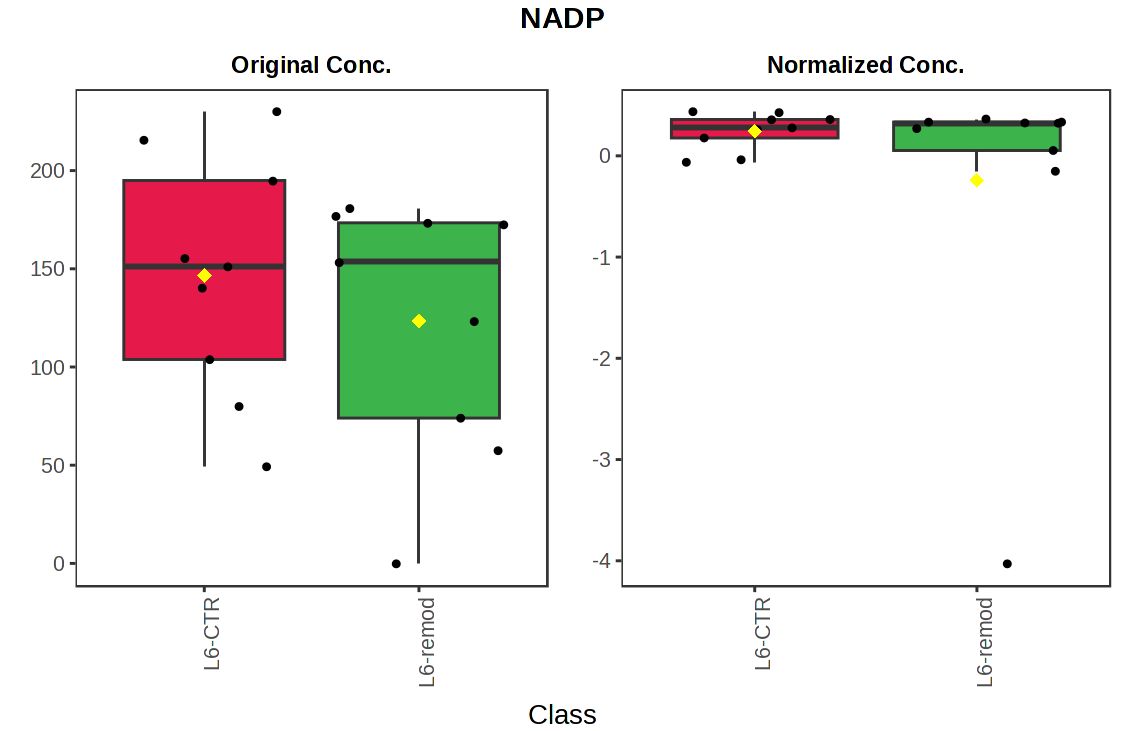

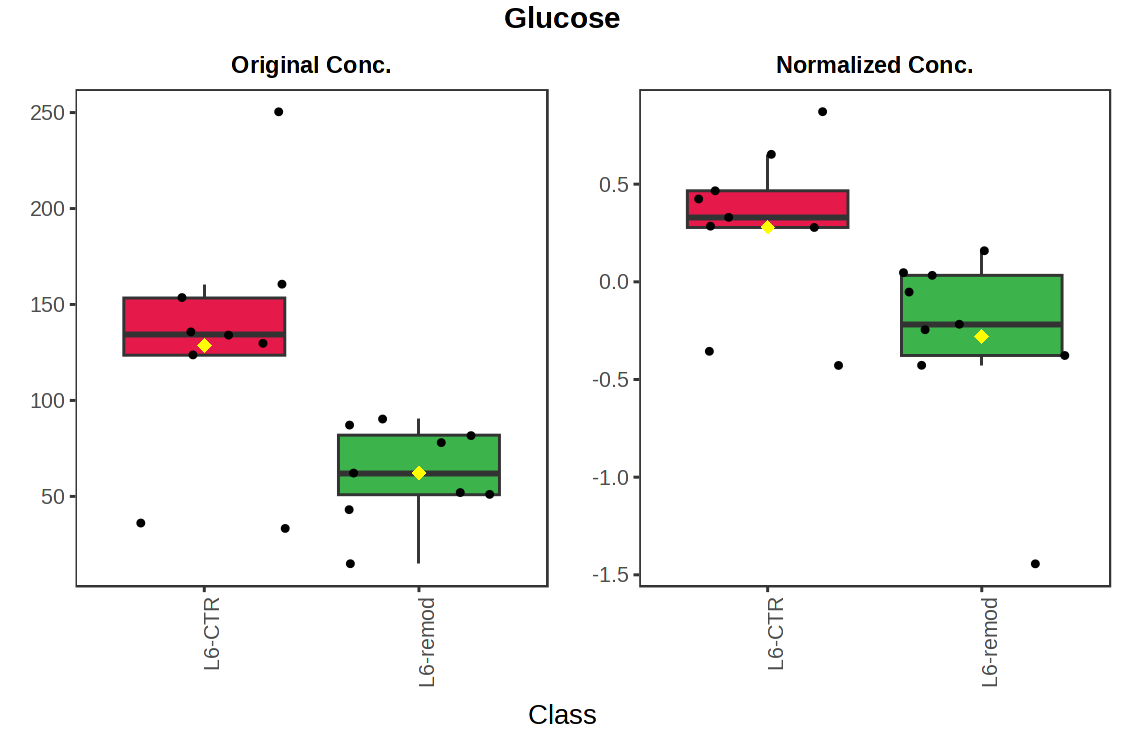

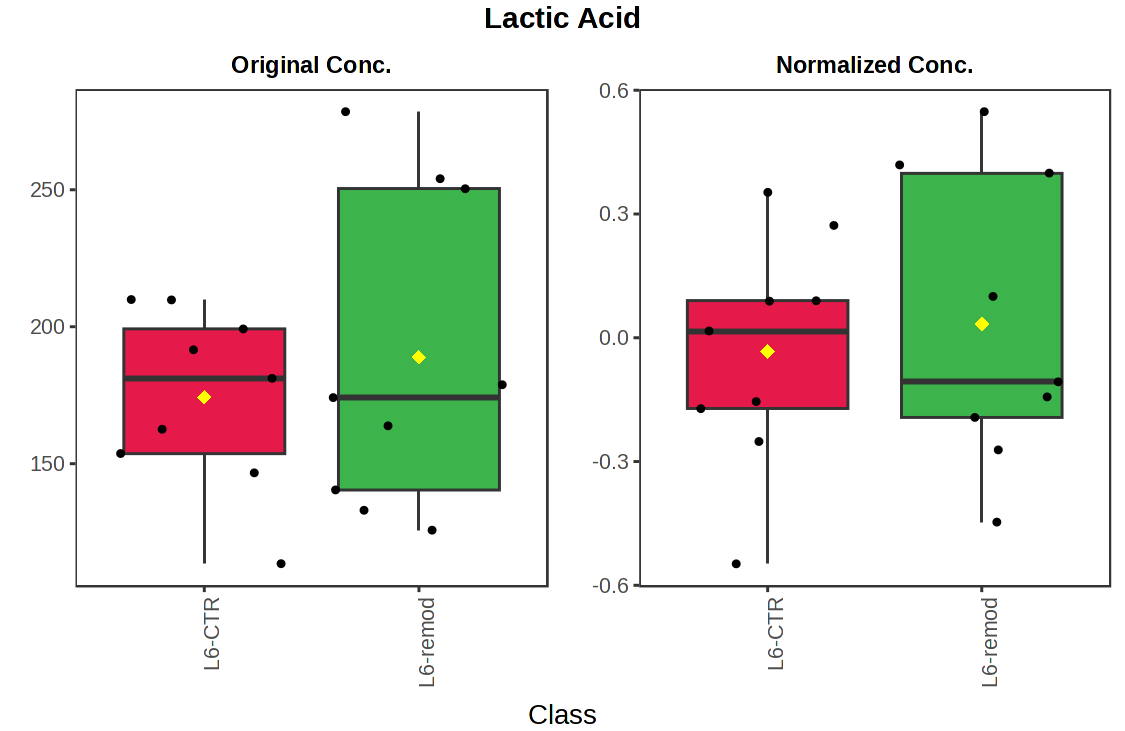

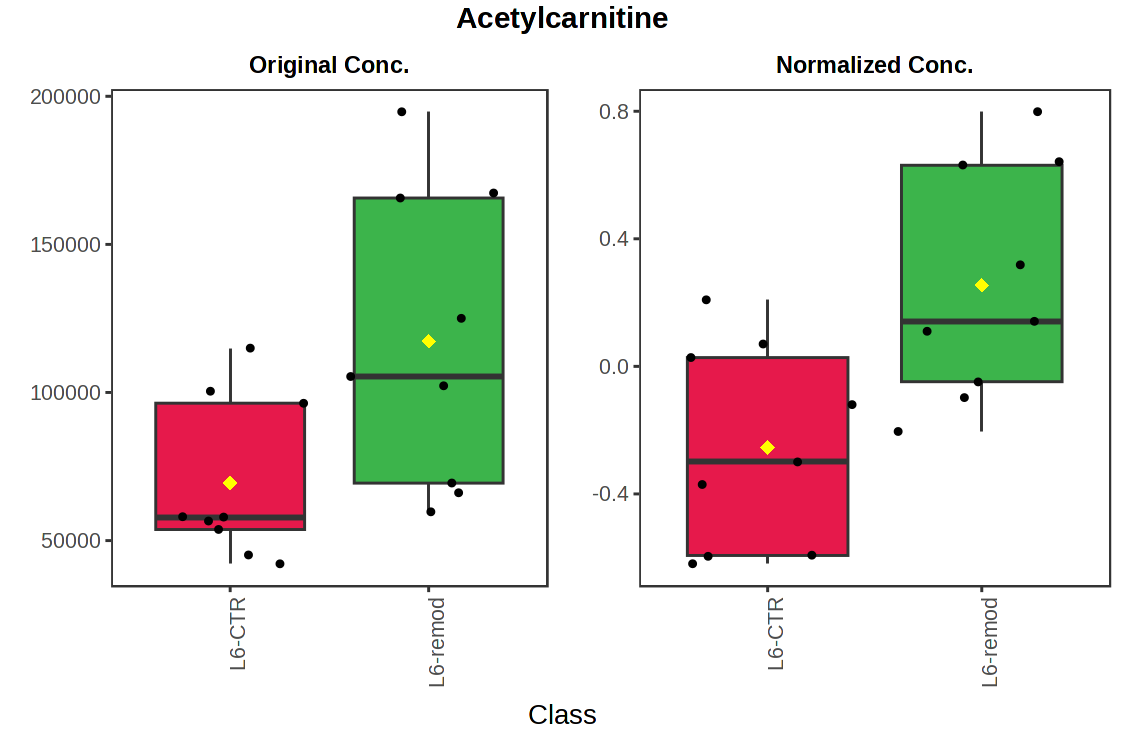

**Acetylcarnitine**

**Lactic acid**

**Glucose**

Normalized Concentration

**Uric Acid**

**NADP**

**UDP Glucuronic Acid**

**L6**

**Control**

**L6**

**Remodelin**

**L6**

**Control**

**L6**

**Remodelin**

**L6**

**Control**

**L6**

**Remodelin**
