## Supplementary Tables for "The inhibitory effects of Remodelin on myoblasts differentiation"

**Supplementary Table S1.**

**C2C12 RNAseq samples**

| **Sample** | **Group of biological replicates** |
| --- | --- |
| M1-M6 | Control day 0 |
| M7-M10 | Control day 3 |
| M11-M14 | Remodelin day 3 |
| M19-M22 | Control day 7 |
| M23-M26 | Remodelin day 7 |
| M31-M34 | Control day 16 |
| M35-M38 | Remodelin day 16 |

**L6 RNAseq samples**

| **Sample** | **Group of biological replicates** |
| --- | --- |
| M1-M4 | Control day 0 |
| M5-M8 | Control day 3 |
| M9-M12 | Control day 7 |
| M13-M16 | Remodelin day 3 |
| M17-M20 | Remodelin day 7 |

**Supplementary Table S2.**

| **Gene Ontology enrichment analysis of downregulated genes at day 3 of differentiation in C2C12 cells** | | | | | |
| --- | --- | --- | --- | --- | --- |
| **Biological process** | | | | | |
| ID | Description | GeneRatio | BgRatio | enrich_factor | pvalue |
| GO:0018101 | protein citrullination | 2.59% | 0.03% | 76.28 | 4,69E-06 |
| GO:0045214 | sarcomere organization | 3.45% | 0.17% | 20.34 | 4,05E-05 |
| GO:0006936 | muscle contraction | 3.45% | 0.2% | 17.54 | 7,42E-05 |
| GO:0046314 | phosphocreatine biosynthetic process | 1.72% | 0.03% | 63.56 | 0,000364 |
| GO:0032924 | activin receptor signaling pathway | 1.72% | 0.06% | 28.25 | 0,00213 |
| GO:1903901 | negative regulation of viral life cycle | 1.72% | 0.07% | 25.43 | 0,002649 |
| GO:0001556 | oocyte maturation | 1.72% | 0.07% | 23.11 | 0,003221 |
| GO:0048525 | negative regulation of viral process | 1.72% | 0.07% | 23.11 | 0,003221 |
| **Cellular component** | | | | | |
| ID | Description | GeneRatio | BgRatio | enrich_factor | pvalue |
| GO:0005883 | neurofilament | 1.57% | 0.05% | 32.62 | 0,001583 |
| GO:0098688 | parallel fiber to Purkinje cell synapse | 1.57% | 0.05% | 29 | 0,002025 |
| GO:0000307 | cyclin-dependent protein kinase holoenzyme complex | 1.57% | 0.07% | 21.75 | 0,003658 |
| GO:0016459 | myosin complex | 2.36% | 0.25% | 9.32 | 0,004058 |
| GO:0000785 | chromatin | 3.15% | 0.54% | 5.86 | 0,004864 |
| GO:0030018 | Z disc | 2.36% | 0.31% | 7.68 | 0,007003 |
| GO:0005615 | extracellular space | 11.02% | 5.3% | 2.08 | 0,007374 |
| GO:0005596 | collagen type XIV trimer | 0.79% | 0.01% | 130.48 | 0,007664 |
| GO:0014801 | longitudinal sarcoplasmic reticulum | 0.79% | 0.01% | 130.48 | 0,007664 |
| GO:0043509 | activin A complex | 0.79% | 0.01% | 130.48 | 0,007664 |
| **Molecular function** | | | | | |
| ID | Description | GeneRatio | BgRatio | enrich_factor | pvalue |
| GO:0004668 | protein-arginine deiminase activity | 2.68% | 0.03% | 86.64 | 3,20E-06 |
| GO:0070080 | titin Z domain binding | 1.79% | 0.01% | 144.4 | 4,75E-05 |
| GO:0003774 | motor activity | 3.57% | 0.24% | 14.81 | 0,000149 |
| GO:0004111 | creatine kinase activity | 1.79% | 0.02% | 72.2 | 0,000283 |
| GO:0051373 | FATZ binding | 1.79% | 0.03% | 57.76 | 0,000469 |
| GO:0036122 | BMP binding | 1.79% | 0.04% | 48.13 | 0,0007 |
| GO:0008083 | growth factor activity | 4.46% | 0.67% | 6.62 | 0,00096 |
| GO:0031432 | titin binding | 1.79% | 0.04% | 41.26 | 0,000976 |
| GO:0003779 | actin binding | 5.36% | 1.13% | 4.76 | 0,00167 |
| GO:0005161 | platelet-derived growth factor receptor binding | 1.79% | 0.07% | 26.25 | 0,00251 |

**Supplementary Table S3.**

| **Gene Ontology enrichment analysis of downregulated genes at day 16 of differentiation in C2C12 cells** | | | | | |
| --- | --- | --- | --- | --- | --- |
| **Biological process** | | | | | |
| ID | Description | GeneRatio | BgRatio | enrich_factor | pvalue |
| GO:0045214 | sarcomere organization | 1.87% | 0.17% | 11.01 | 1,84E-14 |
| GO:0006936 | muscle contraction | 1.98% | 0.2% | 10.09 | 2,26E-14 |
| GO:0006941 | striated muscle contraction | 1.05% | 0.08% | 12.91 | 1,36E-09 |
| GO:0060048 | cardiac muscle contraction | 1.28% | 0.14% | 9.46 | 2,48E-09 |
| GO:0003009 | skeletal muscle contraction | 1.05% | 0.11% | 9.68 | 5,73E-08 |
| GO:0005978 | glycogen biosynthetic process | 0.82% | 0.07% | 12.05 | 2,25E-07 |
| GO:0007517 | muscle organ development | 1.17% | 0.16% | 7.17 | 3,88E-07 |
| GO:0002027 | regulation of heart rate | 0.93% | 0.11% | 8.6 | 1,07E-06 |
| GO:0006096 | glycolytic process | 1.4% | 0.27% | 5.16 | 1,69E-06 |
| GO:0030240 | skeletal muscle thin filament assembly | 0.58% | 0.04% | 14.34 | 3,74E-06 |
| **Cellular component** | | | | | |
| ID | Description | GeneRatio | BgRatio | enrich_factor | pvalue |
| GO:0030018 | Z disc | 2.91% | 0.31% | 9.44 | 7,17E-21 |
| GO:0031430 | M band | 1.18% | 0.09% | 13.08 | 1,80E-11 |
| GO:0030016 | myofibril | 1.61% | 0.19% | 8.63 | 1,96E-11 |
| GO:0030017 | sarcomere | 1.51% | 0.17% | 8.92 | 5,28E-11 |
| GO:0043292 | contractile fiber | 1.4% | 0.18% | 8 | 1,46E-09 |
| GO:0005861 | troponin complex | 0.75% | 0.04% | 17.84 | 1,70E-09 |
| GO:0031674 | I band | 1.18% | 0.13% | 9.34 | 3,41E-09 |
| GO:0005865 | striated muscle thin filament | 0.86% | 0.06% | 14.27 | 3,86E-09 |
| GO:0031672 | A band | 0.86% | 0.07% | 11.89 | 3,83E+06 |
| GO:0044449 | obsolete contractile fiber part | 0.97% | 0.13% | 7.64 | 8,39E-07 |
| **Molecular function** | | | | | |
| ID | Description | GeneRatio | BgRatio | enrich_factor | pvalue |
| GO:0051015 | actin filament binding | 3.33% | 0.79% | 4.21 | 2,13E-12 |
| GO:0008307 | structural constituent of muscle | 1.25% | 0.11% | 11.88 | 8,50E-12 |
| GO:0008017 | microtubule binding | 3.23% | 1.05% | 3.07 | 1,95E-08 |
| GO:0005524 | ATP binding | 12.7% | 8.04% | 1.58 | 2,08E-07 |
| GO:0003779 | actin binding | 3.12% | 1.13% | 2.77 | 3,34E-07 |
| GO:0051373 | FATZ binding | 0.52% | 0.03% | 16.83 | 7,34E-07 |
| GO:0005523 | tropomyosin binding | 0.73% | 0.07% | 9.82 | 1,56E-06 |
| GO:0042802 | identical protein binding | 7.6% | 4.57% | 1.66 | 1,20E-05 |
| GO:0031433 | telethonin binding | 0.42% | 0.02% | 16.83 | 1,24E-05 |
| GO:0051371 | muscle alpha-actinin binding | 0.62% | 0.07% | 9.18 | 1,55E-05 |

**Supplementary Table S4.**

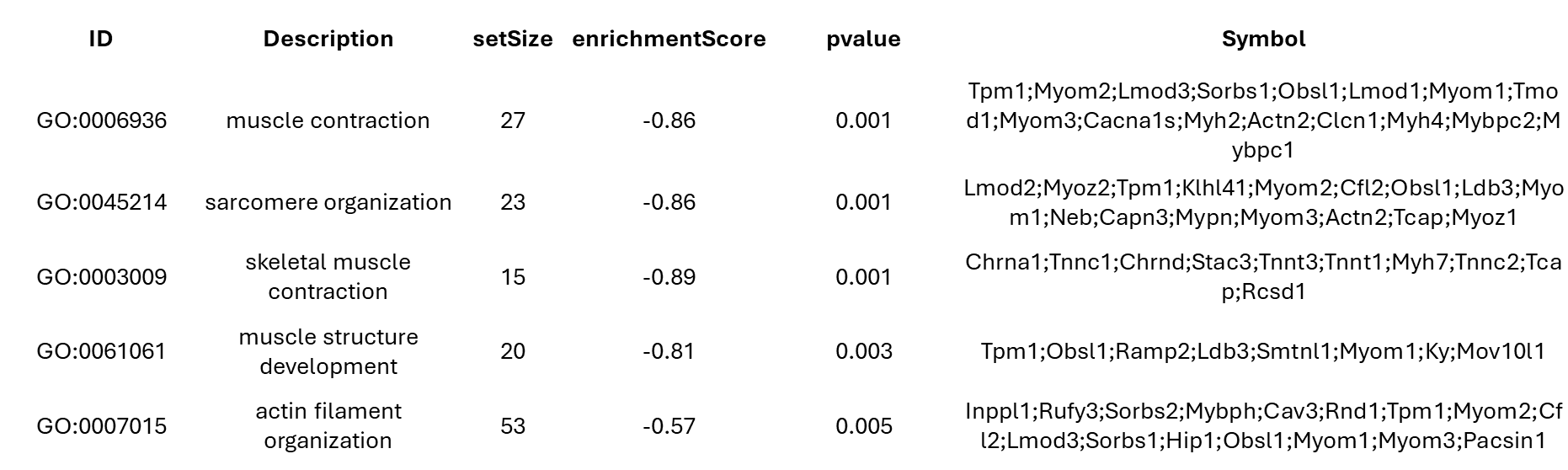

**Supplementary Table S5**

| **Gene Ontology enrichment analysis of downregulated genes at day 7 of differentiation in L6 cells** | | | | | |
| --- | --- | --- | --- | --- | --- |
| **Biological process** | | | | | |
| **term_name** | **term_id** | **adjusted_p_value** | **term_size** | **query_size** | **intersection_size** |
| regulation of biological process | GO:0050789 | 5,89E-12 | 6963 | 625 | 453 |
| cytoskeleton organization | GO:0007010 | 8,75E-08 | 821 | 625 | 111 |
| regulation of actin filament-based process | GO:0032970 | 3,01688E-06 | 207 | 625 | 29 |
| myotube differentiation | GO:0014902 | 6,77093E-06 | 93 | 625 | 18 |
| actomyosin structure organization | GO:0031032 | 7,50389E-06 | 114 | 625 | 20 |
| actin filament organization | GO:0007015 | 2,35518E-05 | 231 | 625 | 29 |
| skeletal muscle tissue development | GO:0007519 | 3,09899E-05 | 137 | 625 | 21 |
| regulation of actin cytoskeleton organization | GO:0032956 | 4,03575E-05 | 187 | 625 | 25 |
| myoblast fusion | GO:0007520 | 0,00013025 | 30 | 625 | 9 |
| **Cellular component** | | | | | |
| **term_name** | **term_id** | **adjusted_p_value** | **term_size** | **query_size** | **intersection_size** |
| cytoskeleton | GO:0005856 | 1,71E-06 | 1324 | 652 | 147 |
| myofibril | GO:0030016 | 6,03E-04 | 120 | 652 | 37 |
| sarcomere | GO:0030017 | 8,59E-04 | 101 | 652 | 34 |
| cytoplasm | GO:0005737 | 7,84E-03 | 6447 | 652 | 412 |
| microtubule organizing center | GO:0005815 | 1,1215E-06 | 593 | 652 | 58 |
| myofilament | GO:0036379 | 3,2652E-05 | 15 | 652 | 7 |
| actin filament bundle | GO:0032432 | 3,31594E-05 | 50 | 652 | 12 |
| troponin complex | GO:0005861 | 5,85137E-05 | 7 | 652 | 5 |
| actomyosin | GO:0042641 | 0,000221071 | 51 | 652 | 11 |
| **Molecular function** | | | | | |
| **term_name** | **term_id** | **adjusted_p_value** | **term_size** | **query_size** | **intersection_size** |
| tubulin binding | GO:0015631 | 2,35726E-06 | 185 | 509 | 30 |
| actin binding | GO:0003779 | 2,86706E-05 | 176 | 509 | 27 |
| extracellular matrix binding | GO:0050840 | 3,62524E-05 | 39 | 509 | 12 |
| laminin binding | GO:0043236 | 0,008208797 | 17 | 509 | 6 |
| tropomyosin binding | GO:0005523 | 0,010039578 | 12 | 509 | 5 |
| cytoskeletal motor activity | GO:0003774 | 0,017442552 | 29 | 509 | 7 |
| actin filament binding | GO:0051015 | 0,029528016 | 84 | 509 | 12 |

| **Supplementary Table S6**  **Gene Ontology enrichment analysis of upregulated genes at day 7 of differentiation in L6 cells** | | | | | |
| --- | --- | --- | --- | --- | --- |
| **Biological process** | | | | | |
| **term_name** | **term_id** | **adjusted_p_value** | **term_size** | **query_size** | **intersection_size** |
| rRNA metabolic process | GO:0016072 | 1,2613E-05 | 76 | 439 | 14 |
| response to oxidative stress | GO:0006979 | 1,62097E-05 | 219 | 439 | 24 |
| response to endoplasmic reticulum stress | GO:0034976 | 2,56768E-05 | 178 | 439 | 21 |
| ribosomal small subunit biogenesis | GO:0042274 | 9,37511E-05 | 45 | 439 | 10 |
| ribosomal large subunit biogenesis | GO:0042273 | 9,37511E-05 | 19 | 439 | 7 |
| lipid metabolic process | GO:0006629 | 0,000108845 | 747 | 439 | 49 |
| regulation of cell adhesion | GO:0030155 | 0,000186287 | 508 | 439 | 37 |
| tRNA modification | GO:0006400 | 0,000753387 | 47 | 439 | 9 |
| nucleolus organization | GO:0007000 | 0,003802998 | 9 | 439 | 4 |
| **Cellular component** | | | | | |
| **term_name** | **term_id** | **adjusted_p_value** | **term_size** | **query_size** | **intersection_size** |
| cytoplasm | GO:0005737 | 2,62E-03 | 6447 | 456 | 310 |
| mitochondrion | GO:0005739 | 4,77E+08 | 926 | 456 | 68 |
| cytosol | GO:0005829 | 9,51E+07 | 2370 | 456 | 129 |
| **Molecular function** | | | | | |
| **term_name** | **term_id** | **adjusted_p_value** | **term_size** | **query_size** | **intersection_size** |
| aminoacyl-tRNA ligase activity | GO:0004812 | 2,4303E-06 | 30 | 392 | 11 |
| protein binding | GO:0005515 | 3,92813E-05 | 4189 | 392 | 211 |
| catalytic activity, acting on RNA | GO:0140098 | 0,000139067 | 228 | 392 | 26 |
| snoRNA binding | GO:0030515 | 0,000368595 | 12 | 392 | 6 |
| ligase activity | GO:0016874 | 0,000937188 | 114 | 392 | 16 |
| transferase activity | GO:0016740 | 0,004283181 | 1505 | 392 | 87 |
| pyridoxal phosphate binding | GO:0030170 | 0,02417127 | 10 | 392 | 4 |
